# Eradication of tumors with pre-existing antigenic heterogeneity by vaccine-mediated co-engagement of CAR T and endogenous T-cells

**DOI:** 10.1101/2022.10.05.511036

**Authors:** Leyuan Ma, Duncan M. Morgan, Ina Sulkaj, Parisa Yousefpour, Charles A. Whittaker, Wuhbet Abraham, Na Li, J. Christopher Love, Darrell J. Irvine

## Abstract

Chimeric Antigen Receptor (CAR) T-cell therapy can be effective in treating human cancer but loss of the antigen recognized by the CAR poses a major obstacle. Here, we report an approach for vaccine boosting CAR T-cells, which triggers engagement of the endogenous immune system to circumvent antigen-negative tumor escape. Vaccine-boosted CAR T-cells promoted dendritic cell (DC) recruitment to tumors, increased tumor antigen uptake by DCs, and elicited priming of endogenous anti-tumor T-cells (antigen spreading). This process was accompanied by a shift in toward oxidative phosphorylation in CAR T-cells and was critically dependent on CAR-T-derived IFN-γ. Antigen spreading induced by vaccine-boosted CAR-T enabled a proportion of complete responses even when the initial tumor was 50% CAR-antigen-negative, and heterogenous tumor control was further enhanced by genetically amplifying CAR-T IFN-γ expression. Thus, CAR T-cell-derived IFN-γ plays a critical role in promoting antigen spreading, and vaccine boosting provides a clinically-translatable strategy to drive such responses against solid tumors.

## INTRODUCTION

Adoptive cell therapy (ACT) using chimeric antigen receptor (CAR) T-cells has revolutionized the treatment of relapsed/refractory CD19^+^ B-cell acute lymphoblastic leukemia and lymphomas (Fesnak et al., 2016; Khalil et al., 2016). In the setting of solid tumors, CAR-T therapy has been less successful so far, though progress is being made in addressing issues such as limited tumor infiltration, poor CAR-T functionality and persistence (Hou et al., 2021; Rafiq et al., 2020; Roselli et al., 2021). However, a key challenge in the treatment of tumors with CAR T-cells is pre-existing antigenic heterogeneity, where not all tumor cells express the antigen targeted by the CAR, or antigen loss occurs during treatment as an adaptive response to immune attack. For example, a recent first-in-human clinical trial assessing CAR T-cells targeting mutant EGFRvIII in glioblastoma resulted in the emergence of EGFRvIII^null^ tumors (O’Rourke et al., 2017). Even in leukemia patients initially responding to CD19 CAR-T therapy, loss or downregulation of the CD19 antigen has been frequently observed and often results in disease relapse (Shah and Fry, 2019). An additional mechanism of antigen loss is via inflammation-induced dedifferentiation in melanomas (Landsberg et al., 2012). These observations highlight the urgent need of novel approaches to address antigen-loss-mediated tumor escape.

Antigen spreading (AS) is the induction and amplification of immune responses to secondary antigens distinct from the original therapeutic target (Gulley et al., 2017). In the setting of adoptive cell therapy, strategies to target one surface-expressed tumor antigen using CAR T-cells while inducing endogenous T-cell responses against additional tumor antigens would be an attractive approach to overcome tumor antigen heterogeneity and antigen loss-mediated escape. Accumulating evidence suggests that AS can be elicited and may contribute to the overall therapeutic outcome during cancer immunotherapy. For example, recruitment and expansion of tumor-specific T-cells that were undetectable prior to therapy was found in patients receiving Ipilimumab (Kvistborg et al., 2014). Some cancer patients treated with neoantigen vaccines also exhibited AS towards overexpressed, shared neoantigens or cancer testis antigens (Awad et al., 2022; Brossart, 2020). However, to date there is limited clinical evidence of CAR T-cell therapy itself inducing therapeutically meaningful AS. In a mesothelin-specific CAR T trial (Beatty et al., 2014), increased antibody responses were observed against tumor-associated antigens as well as weak T-cell responses, but endogenous T-cell clonal expansions were not observed (Kim et al., 2017). A case report of a patient treated with HER2 CAR T-cells for metastatic rhabdomyosarcoma also documented oligoclonal expansion of endogenous T-cells (Hegde et al., 2020), suggesting antigen spreading. Most recently, in a rare case where a GBM patient received local administration of IL13Rα2-targeting CAR T-cells and achieved complete remission, endogenous T-cell responses could be detected during the response (Alizadeh et al., 2021). Overall however, current CAR T therapies appear to be inefficient at promoting endogenous anti-tumor immunity.

Preclinically, a majority of CAR-T studies employ immunodeficient mice that by definition exclude endogenous T-cell responses. When carried out in immunocompetent mouse models, CAR-T therapy itself seems to have limited ability to trigger AS especially in solid tumors (Klampatsa et al., 2020). By contrast, CAR T-cells engineered with additional immune response-provoking molecules, including FLT3L (Lai et al., 2020), CD40L (Kuhn et al., 2020), IL-12 (Etxeberria et al., 2019; Kueberuwa et al., 2018), IL-18 (Chmielewski and Abken, 2017), IL-7/CCL19 (Adachi et al., 2018), or when used in combination with oncolytic viruses (Park et al., 2020; Walsh et al., 2019), have been reported to exhibit increased anti-tumor activity as well as evidence for AS. However, introduction of such additional effector functions with uniform activity across patients can be challenging and introduces new safety risks (Kerkar et al., 2010; Zhang et al., 2015). More importantly, irrespective of the CAR T-cell modality, mechanisms by which AS is promoted during adoptive cell therapy remain poorly understood.

We recently described an approach to amplify CAR-T activity in solid tumors by vaccine-like boosting of CAR T-cells via their chimeric antigen receptor in lymph nodes (Ma et al., 2019). This was accomplished by the synthesis of CAR ligands conjugated to an amphiphilic polymer-lipid tail, which following parenteral injection, efficiently traffic to draining lymph nodes and decorate the surfaces of macrophages and dendritic cells (DCs) with CAR T ligands. CAR T-cells encountering ligand-decorated DCs in the lymph node receive stimulation through the CAR in tandem with native costimulatory receptor signals and cytokine stimulation from the ligand-presenting cell, leading to CAR T-cell expansion and enhanced functionality. Vaccine boosting of CAR T-cells via administration of these “amph-ligands” together with vaccine adjuvants substantially enhanced tumor rejection by CAR T-cell therapy, and unexpectedly, was accompanied by the development of endogenous anti-tumor T-cell responses (Ma et al., 2019).

Here we used this approach of CAR-T therapy in tandem with vaccine boosting as a model setting to understand the role of antigen spreading in effective clearance of antigenically heterogenous solid tumors, and to define mechanisms underlying antigen spreading induced by CAR T-cell attack on solid tumors. In multiple murine syngeneic tumor models, we found that AS elicited by CAR T-cell treatment using second-generation CARs currently used in the clinic was negligible. However, endogenous T-cell priming could be markedly induced by vaccine boosting of CAR T-cells, even in the context of lymphodepletion preconditioning. This process was critically dependent on IFN-γ, involving both autocrine stimulation of CAR T-cells and paracrine stimulation of the host immune system, reinforced by host IL-12 signaling and at least partially sustained by vaccine-mediated reprogramming of CAR T-cell metabolism. Importantly, enhanced IFN-γ expression induced either by vaccine boosting or genetic engineering enabled CAR T-cells to control solid tumors with preexisting antigen heterogeneity.

## RESULTS

### Vaccine boosting enables CAR T-cells to elicit endogenous CD4^+^ and CD8^+^ T-cell responses in multiple tumor models

Steps in the amph-ligand-based vaccine boosting approach are illustrated schematically in **Figure 1A**: Amph-ligands are comprised of a ligand for a selected CAR linked to a hydrophobic phospholipid tail via a poly(ethylene glycol) (PEG) spacer. Upon co-injection with a suitable vaccine adjuvant at a site distal from the tumor, amph-ligands bind to albumin present in the interstitial fluid and are efficiently transported to the downstream draining lymph nodes (dLNs) (Liu et al., 2014). Within the densely packed LN parenchyma, the amph-ligand transfers into cell membranes, decorating primarily the surface of macrophages and dendritic cells that line the subcapsular sinus and collagen conduits carrying lymph into the T-cell paracortex (Ma et al., 2019). The co-administered adjuvant simultaneously activates DCs in the dLN to upregulate expression of costimulatory receptors and produce cytokines. CAR T-cells encountering ligand-decorated, activated DCs are stimulated in a manner mimicking natural T-cell priming, leading to CAR T-cell expansion and enhanced effector functions. Unexpectedly, we found that vaccine-boosted CAR T-cells also induce the expansion of endogenous anti-tumor T-cell responses (Ma et al., 2019).

**Figure. 1.**
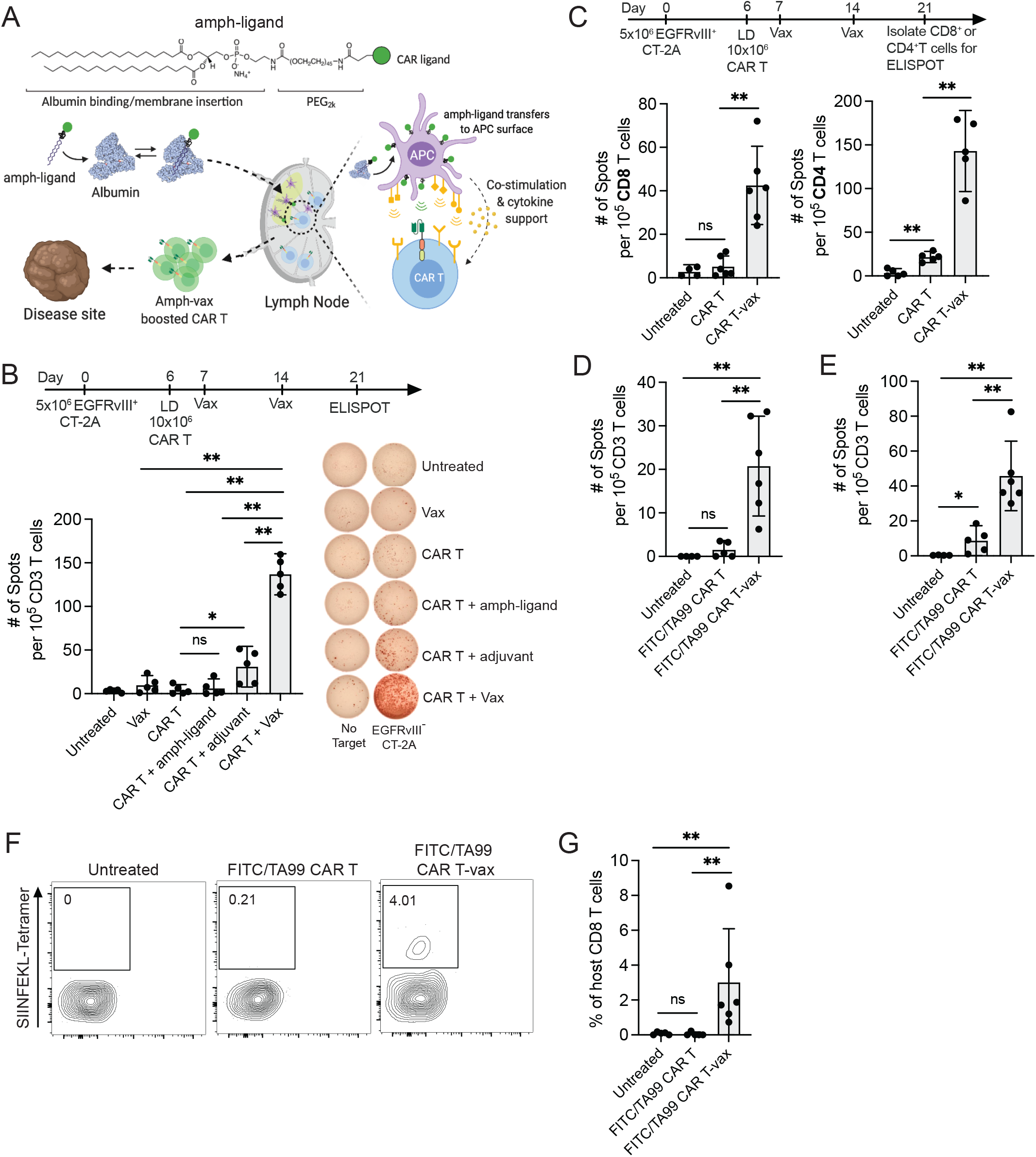
Vaccine boosting enables CAR T-cells to elicit endogenous T-cell responses in multiple tumor models. (A) Schematic of CAR T-vax therapy. (B) C57BL/6 mice bearing EGFRvIII^+^CT-2A tumors received lymphodepletion (LD) and were left untreated or treated with CAR T + various combination of vaccine components as shown in the timeline. Shown is IFN-γ ELISPOT monitoring endogenous T-cell priming across various conditions. Shown is one representative of three independent experiments. (C) Priming of endogenous CD4 and CD8 T-cell in CT-2A tumor-bearing mice upon treatment with or without CAR T ± vax as measured by IFN-γ ELISPOT following T cell restimulation with irradiated EGFRvIII^-^ CT-2A cells. Shown above is the timeline. (D-G) C57BL/6 mice (n=5-6 animals/group) bearing OVA^+^ B16F10 tumors received FITC/TA99 CAR T-vax treatment as in Fig 1C, and T cells were isolated from splenocytes on day 21 for analysis. Shown is one representative of two independent experiments. (D) IFN-γ ELISPOT measuring OVA-specific endogenous T-cell responses following restimulation with irradiated OVA-expressing CT-2A cells. (E) IFN-γ ELISPOT measuring endogenous T-cell responses following restimulation with irradiated Trp1^-/-^ B16F10 cells. (F-G) Tetramer-staining showing representative flow cytometry staining (F) and mean percentages of SIINFEKL tetramer^+^ endogenous T cells (G). Error bars show mean ± 95% CI. ***, p<0.0001; **, p<0.01; *, p<0.05; n.s., not significant by one-way ANOVA with Tukey’s post-test.

We first assessed how the composition of the boosting vaccine impacts this antigen spreading response in a syngeneic murine EGFRvIII^+^ CT-2A glioblastoma model. In this model, CAR T-cells targeting mutant EGFR (mEGFRvIII) are vaccine boosted using an amph-ligand comprised of an mEGFRvIII-derived peptide epitope recognized by the CAR T-cells (Ma et al., 2019) (**Figure S1A**) combined with the potent STING agonist vaccine adjuvant cyclic di-GMP. Mirroring clinical protocols, we lymphodepleted animals prior to adoptive transfer to promote CAR T-cell engraftment, followed one day later by s.c. injection of amph-ligand alone, adjuvant alone, or the full vaccine (amph-ligand + adjuvant). Amph-ligand/adjuvant was administered again 7 days later as a second boost, and then splenocytes were isolated at day 21 and co-cultured with irradiated EGFRvIII^-^ CT-2A cells in an IFN-γ ELISPOT assay to detect endogenous T-cell responses against non-CAR-T-targeted antigens (**Figure 1B**). Endogenous lymphocyte and dendritic cell numbers were still recovering across the time course of these experiments following lymphodepletion (**Figure S1B**). However, their recovery was sufficiently rapid to permit robust *de novo* endogenous T-cell priming, consistent with prior preclinical studies reporting antigen spreading following lymphodepleting therapy (Lai et al., 2020). Injection of the amph-ligand alone without adjuvant failed to elicit endogenous T-cell priming, while CAR T treatment in tandem with vaccine adjuvant alone could promote a low but detectable endogenous T-cell response (**Figure 1B**). However, the full vaccine (amph-ligand + adjuvant) led to 6-fold greater endogenous T-cell priming. This antigen spreading response was not a result of any direct effect of the vaccine on tumors, as inoculating tumors distal from the vaccine injection site did not change the antigen spreading response (**Figure S1C**). Similar magnitudes of endogenous T-cell priming were also observed with alternative adjuvants (TLR7/8 agonist Resiquimod, or the TLR9 agonist CpG, **Figure S1D**), suggesting that a variety of known vaccine adjuvants can be effective in promoting CAR T-cell-driven AS. Isolation of CD4^+^ or CD8^+^ T-cells from the spleens of mice bearing EGFRvIII^+^ CT-2A tumors following treatment revealed that while CAR-T therapy elicited no statistically significant endogenous anti-tumor CD8^+^ T-cell response and only a weak (but detectable) CD4^+^ T-cell response compared to untreated tumors, CAR T combined with amph-ligand vaccination (hereafter, CAR T-vax) primed robust responses from both the CD4^+^ and CD8^+^ T-cell compartments (**Figure 1C**).

To explore the generality of this approach for promoting endogenous anti-tumor immunity and to evaluate AS in a tumor model carrying a defined T-cell antigen, we assessed vaccine-boosted CAR T treatment in a second model of B16F10 murine melanoma expressing the surrogate antigen ovalbumin (OVA), treated with bispecific CAR T-cells recognizing the melanoma-associated antigen Trp1 and FITC (**Figure S1E**). In this model, the CAR T-cells are boosted by vaccination with amph-FITC and attack the tumor through Trp1 recognition. By ELISPOT, we observed endogenous T-cell responses to both the model antigen OVA (**Figure 1D**) and B16F10 neoantigens (**Figure 1E**), but only when mice received both CAR T-cells and vaccine boosting. As a complementary approach to measure endogenous responses to a tumor antigen irrespective of their particular effector function, we also treated B16F10-OVA tumors with CAR T-cells ± vaccine boosting and quantified CD8^+^ T-cells targeting the immunodominant OVA epitope SIINFEKL by peptide-MHC tetramer staining. As shown in **Figure 1F-G**, no OVA-specific T-cells were detected in untreated tumor-bearing animals or mice receiving CAR T-cells alone, but CAR T treatment in the presence of vaccine boosting elicited a readily detectable SIINFEKL-specific T-cell response. Thus, in two different tumor models using two different CARs, CAR T-cell treatment combined with amph-vax boosting promoted endogenous anti-tumor T-cell priming.

### Vaccine-boosted CAR T-cells drive functional enhancement and phenotypic changes in endogenous T-cells

We next sought to characterize the roles of CAR T-vax treatment on the functional state of the endogenous T-cell response. As endogenous tumor-specific T-cells primed by CAR T-vax treatment would be expected to infiltrate the tumor, we analyzed endogenous tumor-infiltrating lymphocytes (TILs) by flow cytometry on day 7 post treatment, and observed substantially increased CD8^+^ TILs and a trend toward increased CD4^+^ cells (**Figure 2A**). Similar increased endogenous T-cell infiltration was found by adding vaccine boosting to CAR-T therapy treatment of OVA-expressing CT2A tumors, and this included a 3-fold increase in *bona fide* tumor-antigen (OVA)-specific TILs (**Figure S1F**). We next isolated host CD45.2^+^ CD4^+^ and CD8^+^ tumor-infiltrating lymphocytes (TILs) 7 or 14 days after initiation of therapy using CD45.1^+^ CAR T-cells, and carried out single-cell RNA-seq and paired α/β TCR sequencing on the recovered host lymphocytes (**Figure 2B**). We obtained quality single-cell transcriptomes for 21,835 T-cells (**Figure 2C-D**). Unsupervised clustering of the transcriptome data revealed five major endogenous T-cell subsets: CD8^+^ cytotoxic T lymphocytes (CTLs, expressing *Cd8a, Ccl5, Pdcd1),* CD4^+^ T helper cells (*Cd4, Cd40lg*), Tregs (*Foxp3, Il2ra, Ikzf2*), a proliferating Ki-67^+^ population that included both CD4^+^ and CD8^+^ cells (*Mki67, Top2a*), and a small population of IFN-stimulated T-cells (characterized by expression of *Ifit1, Ifit3, Isg15)* (**Figure 2D-E, S2A, Supplemental Table 1**), as has been described previously (Szabo et al., 2019; Tibbitt et al., 2019). We observed an increase in the frequency of the CD8^+^ CTL population in mice treated with CAR T-vax at both day 7 and day 14. We also observed a decrease in the frequency of Tregs at day 7 in mice treated with CAR T-vax compared to those treated with CAR-T alone (**Figure 2E**).

**Figure. 2.**
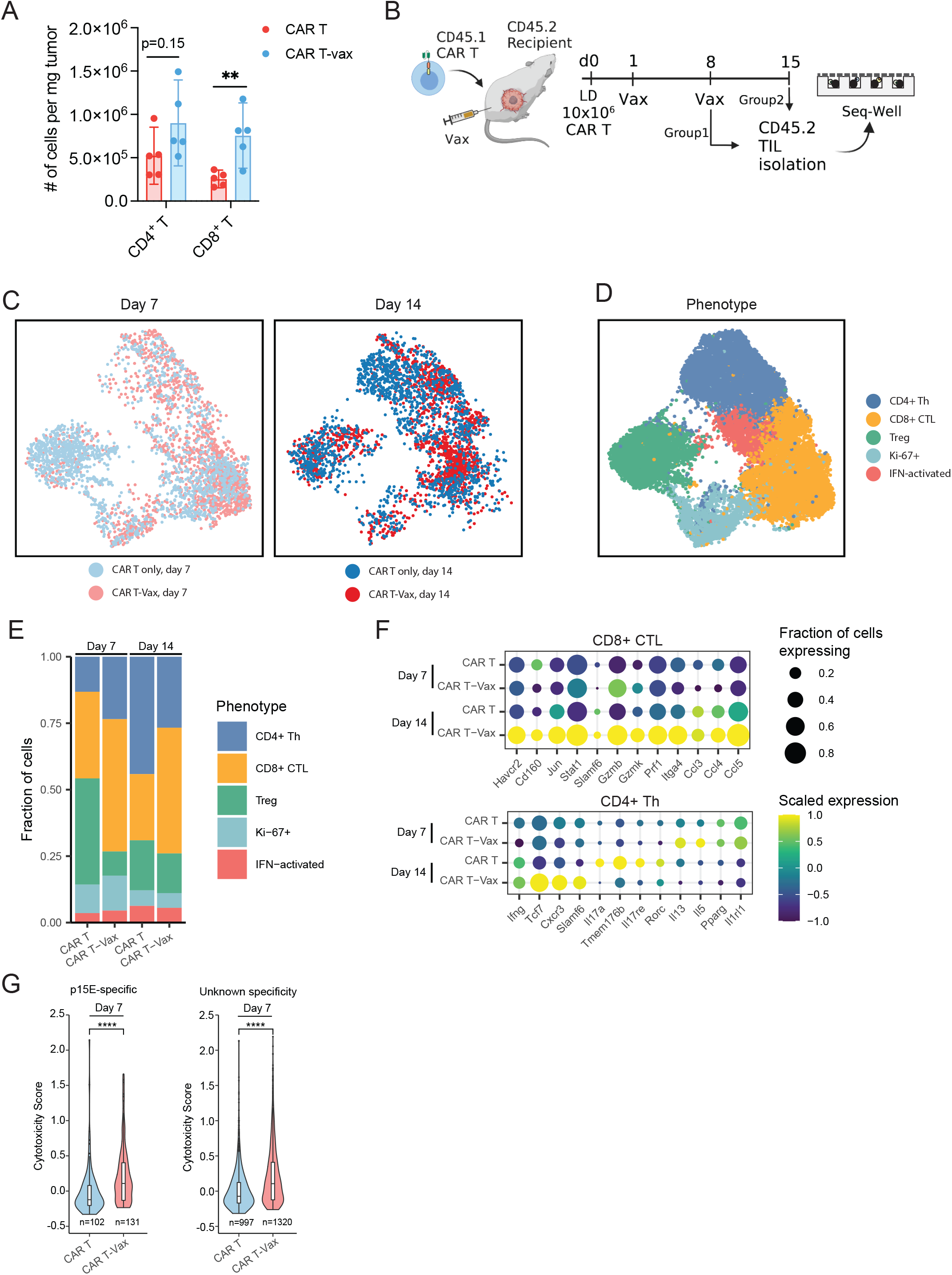
Endogenous tumor-infiltrating T cells show transcriptional changes associated with enhanced anti-tumor activity in response to CAR T-vax therapy. (A) Enumeration of intratumoral CD4^+^ and CD8^+^ host T-cells in mice bearing CT-2A tumors (n=5 animals/group) receiving CAR T ± vax treatment as in Fig. 1C. Error bars show mean ± 95% CI. **, p<0.01; n.s., not significant by Student’s t-test. (B-G) CD45.2^+^ mice bearing CT-2A tumors were treated with EGFRvIII-targeting CD45.1+ CAR T-cells with or without vaccine boosting, followed by TIL isolation on day 7 or day 14 post-vaccination for scRNA-seq analysis using Seq-Well. (B) Experimental setup and timeline (C) UMAP of endogenous T-cells obtained from tumors on day 7 or day 14, colored by the treatment group. 21,835 cells were sequenced; plot shows data randomly down-sampled to show an even number of points from each treatment condition. (D) Curated clusters based on signature gene expression. Th, T helper cells. Treg, regulatory T-cell. CTL, cytotoxic lymphocyte. (E) Stacked charts showing proportions of each T-cell cluster present following each treatment condition on day 7 and day 14. (F) Dot plots showing differential expression of signature genes in endogenous CD8^+^ CTLs or CD4^+^ Th cells following CAR T-vax or CAR-T alone treatment on day 7 and day 14. (G) Cytotoxicity score of endogenous retroviral antigen p15E-specific TILs and TILs of unknown specificity on day 7 for each treatment condition. Error bars are mean ± 95% CI, ****p<0.0001, by two-sided Wilcoxon rank-sum test. See methods for the definition and calculation of the cytotoxicity score.

To assess whether CAR T-vax is able to alter the phenotype of endogenous CD8^+^ CTLs in the tumor microenvironment, we computed differentially expressed genes between CD8^+^ CTLs recovered from mice treated with CAR T-vax vs. CAR-T alone. At day 14, CD8^+^ T-cells from CAR T-vax-treated mice upregulated transcripts associated with both cytotoxicity (*Gzmb, Gzmk)* and T-cell activation (*Havcr2*) relative to cells in the CAR-T alone group (**Figure 2F, S2B-C, Supplemental Table 2**). We validated these findings at the protein level by carrying out flow cytometry analysis of endogenous TILs. Compared to CAR-T only therapy, vaccine boosting did not change the proportion of PD-1^+^TIM-3^+^ or PD-1^+^TIM-3^-^ endogenous TILs (**Figure S3A**), but did enhance IFN-γ, TNF-α, and granzyme B expression in both of these populations (**Figure S3B-C**). Among CD4^+^ cells, we found an elevation of transcripts associated with Th17 function (*Rorc, Il17a, Il17re*) among mice treated with CAR-T alone at day 14 compared to day 7 (**Figure 2F, Supplemental Table 2**). By contrast, CD4^+^ Th cells from mice treated with CAR T-vax upregulated genes associated with Th1 function (*Ifng, Cxcr3*) and self-renewal (*Slamf6, Tcf7*) (**Figure 2F, S2D-E**), suggesting that the vaccine may also promote anti-tumor phenotypes among CD4^+^ TILs. Next we sought to assess how CAR T-vax affects TILs according to their antigen specificities. Using data generated in a recent study defining TCR sequences specific for a common murine endogenous retroviral antigen p15E (Grace et al., 2022) that is also expressed by CT-2A cells (**Figure S3D-E**), we assessed the transcriptional state of tumor-specific endogenous TILs. At day 7, both p15E-specific T-cells and TILs of unknown specificity from CAR T-vax-treated mice exhibited significantly higher cytotoxicity than TILs from animals treated with CAR-T alone (**Figure 2G, S2 F-G, S3F, Supplemental Table 3**). Overall, this analysis suggests that the addition of the vaccine to the CAR T therapy regimen increases the anti-tumor potential of tumor-infiltrating CD8^+^ T-cells and skews the differentiation of tumor-infiltrating CD4^+^ T-cells to a Th1 phenotype.

### Vaccine-driven antigen spreading prevents relapse of antigen-loss variants and enables control of antigenically heterogenous tumors

To determine if endogenous T-cells expanded by CAR T-vax therapy impact the outcome of treatment, we treated wild-type (WT) or RAG1^-/-^ mice bearing mEGFRvIII^+^CT-2A tumors with CAR T-cells ± vaccine boosting. Early tumor regression was observed following CAR T-vax treatment in both WT and RAG1^-/-^ mice, but a majority of tumors in RAG^-/-^ animals relapsed 20-50 days post treatment, even in cases where the tumor appeared to be fully rejected (**Figure 3A**). By contrast, tumors that regressed and reached an undetectable size following CAR T-vax treatment became long-term survivors with no relapses for at least 105 days (**Figure 3A-B**). Flow cytometry analysis of EGFRvIII expression on relapsed tumors revealed that loss or down-regulation of EGFRvIII on tumor cells was a major escape mechanism in the RAG^-/-^ animals (**Figure 3C**).

**Figure. 3.**
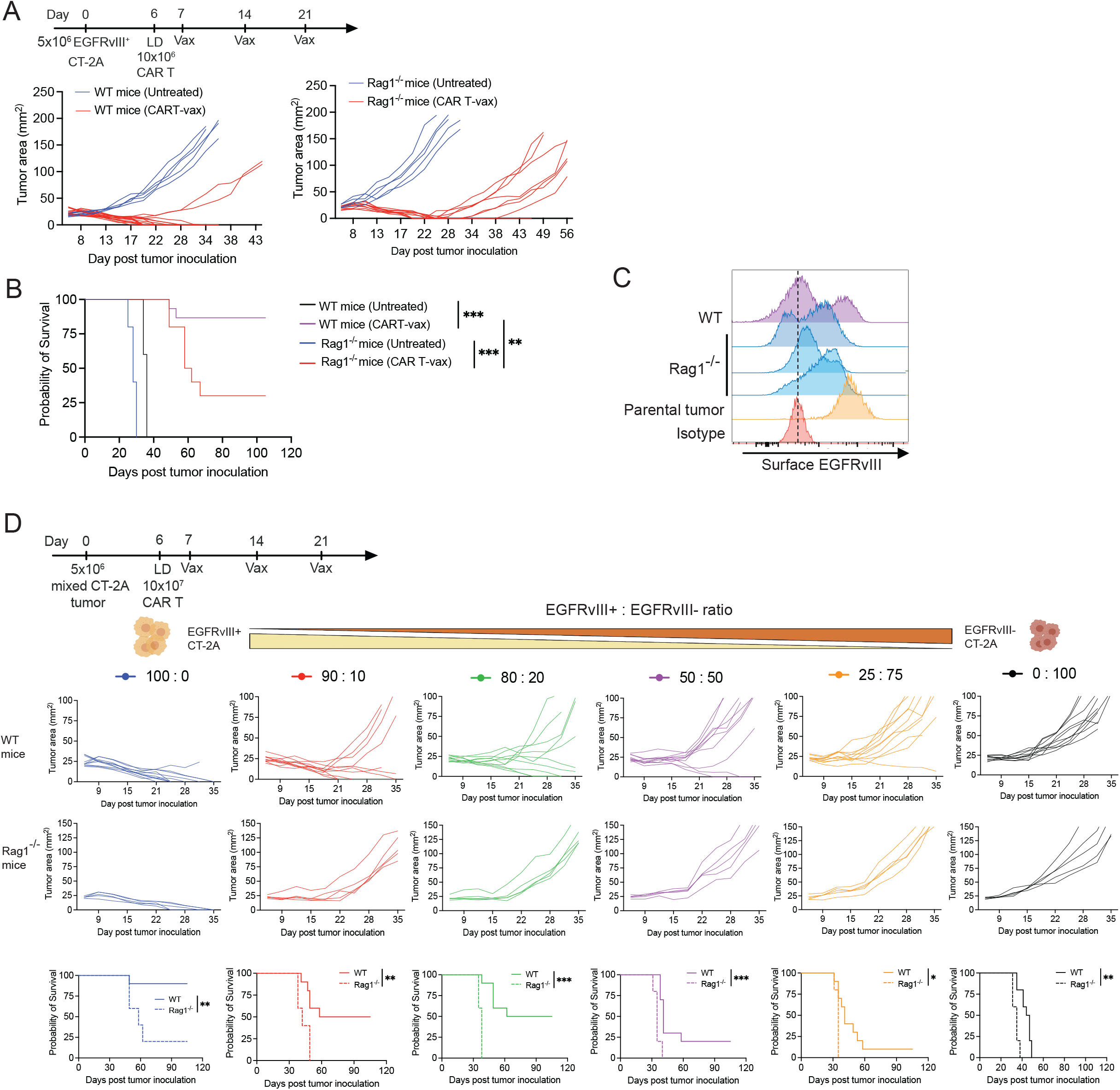
Vaccine-driven antigen spreading is required for long-tumor tumor control in immunocompetent mice. (A-C) EGFRvIII-CT-2A tumor-bearing mice were left untreated or received EGFRvIII-CAR T-vax therapy in wild type (WT, *n*=10 animals/group) or Rag1^-/-^ mice (*n*=5 animals/group). Timeline of experiment is shown above panel (A). Shown is the combined data from two independent experiments. (A) Tumor growth in individual mice. (B) Overall survival. (C) Surface EGFRvIII expression on parental or representative relapsed tumors collected on day 35 from WT and Rag1^-/-^ mice. (D) Mixtures of EGFRvIII^+^ CT-2A and EGFRvIII^-^ CT-2A tumor cells at the indicated ratios were inoculated in WT (*n*=10 animals/group) or Rag1^-/-^ mice (*n*=5 animals/group), followed by lymphodepletion and treatment with CAR T-vax therapy as shown in the timeline. Shown are individual tumor growth curves and overall survival. Shown in WT mice is the combined data from two independent experiments. In panels B and D, ***, p<0.0001; **, p<0.01; *, p<0.05 by Log-rank (Mantel-Cox) test.

Encouraged by these findings, we tested whether endogenous T-cell priming could enable CAR T-cells to eliminate tumors with pre-existing antigenic heterogeneity. We established a heterogeneous tumor model by inoculating a mixture of EGFRvIII^+^ CT-2A cells and parental EGFRvIII^-^ CT-2A cells at defined ratios into both WT and RAG1^-/-^ mice (**Figure 3D**). We previously showed that these two CT-2A variants have similar growth rates in WT mice (Ma et al., 2019). When 100% of the tumor cells express EGFRvIII, CAR T-vax therapy elicited comparable initial tumor regressions in both WT and RAG1^-/-^ mice, but long-term remission was only achieved in WT mice (**Figure S3G**). More strikingly, in heterogeneous tumors comprised of as little as 10% EGFRvIII^-^ cells, CAR T-vax therapy delayed tumor progression but induced no actual regressions in RAG1^-/-^ mice. By contrast, CAR T-vax treatment cured ~50% animals bearing tumors with up to 20% EGFRvIII^-^ cells, and could still achieve complete responses in a small proportion of animals when the EGFRvIII^-^ population was 50% of the tumor mass at time zero. This drastic difference of therapeutic outcome in WT vs RAG1^-/-^ mice demonstrates the pivotal role endogenous T-cells and AS can play in controlling tumors with pre-existing antigenic heterogeneity.

### Vaccine boosting induces cell-intrinsic enhancements in CAR T-cell function

We next sought to understand how amph-vax boosting promoted endogenous T-cell priming. We first evaluated whether the anti-tumor efficacy of CAR T combined with vaccination was simply driven by increased numbers of CAR T-cells induced by vaccination, vs. a change in CAR-T function. To distinguish these possibilities, CAR T-cells were transferred into non-tumor bearing mice, vaccine boosted (or not as controls), and then isolated 7 days later from the two groups and transferred at equal numbers into new tumor-bearing recipient mice (**Figure 4A**). This approach revealed that even when the same number of CAR T-cells were present, vaccine-boosted CAR-T still exhibited enhanced tumor control and long-term animal survival, suggesting that vaccine boosting enhances the intrinsic per-cell functionality of CAR T-cells (**Figure 4A**).

**Figure. 4.**
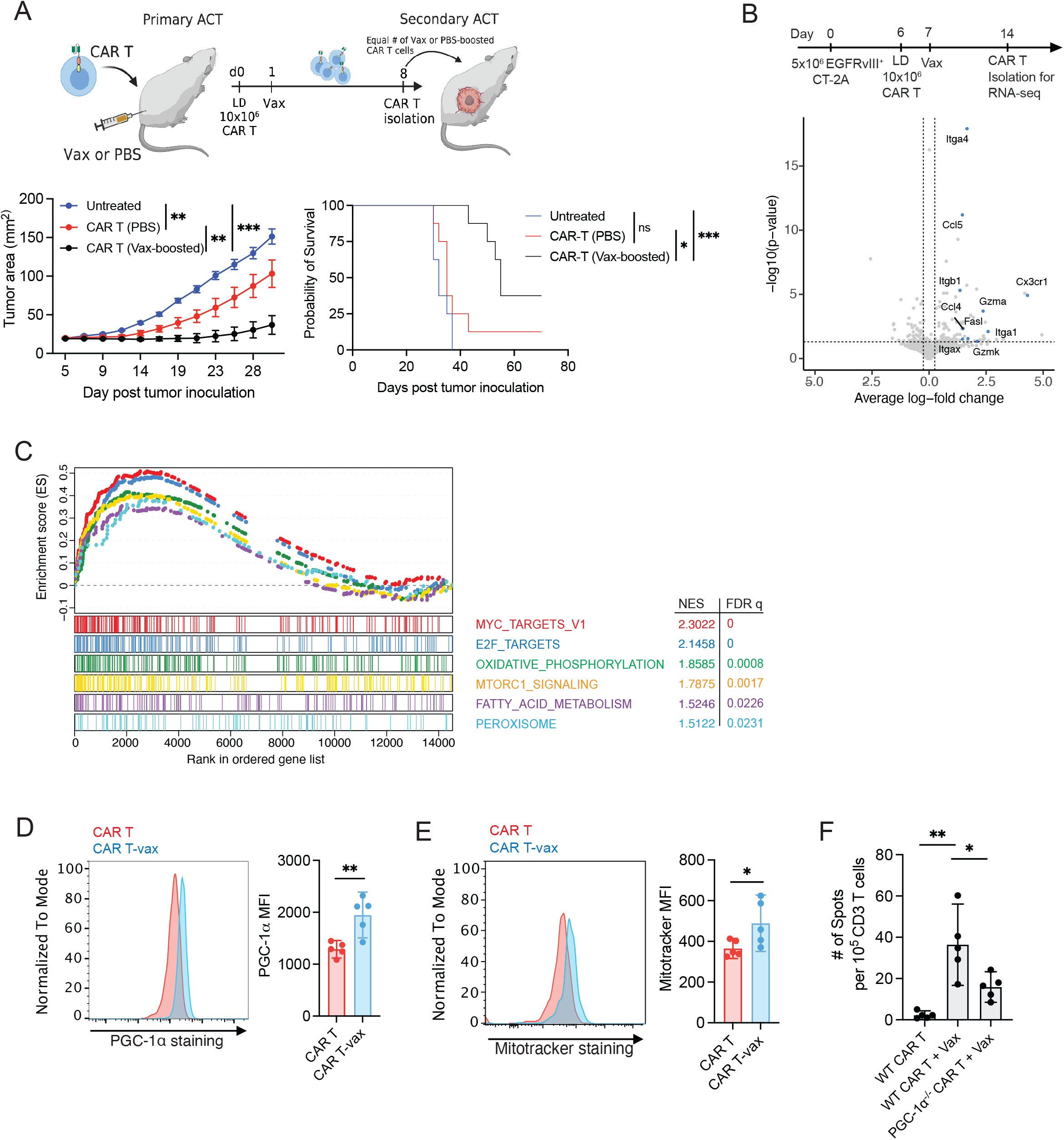
Vaccine boosting induces cell-intrinsic enhancements in CAR T-cell function that include metabolic reprogramming. (A) CAR-T cells were transferred into recipient mice and vaccine-boosted or not, and 7 days later equivalent numbers of boosted or non-boosted cells were recovered and transferred into secondary CT-2A tumor-bearing hosts (*n* = 8 animals/group). Shown are average tumor growth (left) and overall survival (right). ns, not significant; *, p < 0.05; **, p < 0.01; ***, p < 0.001 by two-way ANOVA with Tukey’s post-test (for tumor growth) or Log-rank (Mantel-Cox) test (overall survival). (B-C) CT-2A tumor-bearing mice were treated with CAR T-cells ± vaccine boosting, and CAR T-cells were isolated on day 14 from spleens and tumors for RNA-seq. (Timeline shown in B). (B) Volcano plot showing differential gene expression in splenic CAR T-cells in the presence or absence of vaccine boosting. Highlighted genes are significantly upregulated in CAR T-cells by vaccine boosting. (C) Gene set enrichment analysis (GSEA) showing highly enriched pathways in intratumoral CAR T-cells upon vaccine boosting. (D-E) CT-2A tumor-bearing mice (n = 5 animals/group) were treated with EGFRvIII-CAR T-cells as in (B) ± vaccine boosting, and on day 7, CAR T-cells were recovered from tumor for flow cytometry analysis of intracellular PGC-1α expression (D) and mitochondrial mass (E). Shown in WT mice is one representative of two independent experiments. (F) CT-2A tumor-bearing mice (n = 5 animals/group) were treated with wild type (WT) or PGC-1 α^-/-^ CAR T-cells as in (B) ± vaccine boosting. On day 21, endogenous T-cell responses in the spleen were assayed by restimulation of T cells in an IFN-γ ELISPOT assay with irradiated EGFRvIII^-^ CT-2A cells. In panel D-F, n=5 animals/group, data shows mean ± 95% CI, **, p<0.01; *, p<0.05 by Student’s *t*-test for D-E, and one-way ANOVA with Tukey’s post-test for F.

To gain an unbiased view of how vaccination alters CAR-T function, we carried out bulk RNA-seq on CAR T-cells that did or did not receive vaccine boosting 8 days after adoptive transfer. Vaccination significantly increased the expression of genes associated with effector function (e.g, *FasL, GZMA, GZMK*) in CAR T-cells recovered from the spleen (**Figure 4B**, **Supplemental Table 4**). Gene set enrichment analysis (GSEA) on tumor-infiltrating cells further revealed that vaccine-boosted CAR T-cells maintained a high proliferative potential, as evidenced by elevated Myc and E2F target genes, suggesting resistance to the immunosuppressive TME (**Figure 4C**). Vaccine-boosted cells also showed a significant upregulation of major metabolic signaling pathways, including oxidative phosphorylation (OXPHOS), MTORC1 signaling, fatty acid metabolism, and peroxisome signaling (**Figure 4C**). Prompted by these transcriptional signatures, we analyzed the intracellular expression of PGC-1α, a master transcription factor controlling many genes and pathways involved in OXPHOS (LeBleu et al., 2014), and found that vaccine boosting increased PGC-1α levels in CAR T-cells (**Figure 4D**). PGC-1α is involved in mitochondria generation and maintenance (Fernandez-Marcos and Auwerx, 2011), and we also noted increased mitochondrial mass in vaccine-boosted CAR T-cells (**Figure 4E**). Notably, endogenous T-cell priming was significantly reduced ~50% following CAR T-vax treatment with PGC-1α^-/-^ CAR T-cells compared to WT CAR-T (**Figure 4F**). Hence, metabolic reprogramming in vaccine-boosted CAR T-cells is one factor promoting antigen spreading.

### Enhanced IFN-γ production by vaccine-boosted CAR T-cells is critical for induction of antigen spreading

OXPHOS has been shown to be critical for maintaining the polyfunctionality of T-cells within the TME (Vardhana et al., 2020), and we previously observed that vaccine-boosted CAR T-cells recovered from the peripheral blood showed increased cytokine production (Ma et al., 2019). To determine if this enhanced effector function was maintained in tumors and impacted antigen spreading, we first assessed production of IFN-γ and TNF-α by tumor-infiltrating CAR T-cells. Expression of both cytokines was markedly increased for CAR T that received vaccine boosting (**Figure 5A**). Interestingly, although boosted/non-boosted CAR T-cells exhibited comparable levels of PD-1 and Tim-3 expression, high-level cytokine production was maintained in both PD-1^+^Tim-3^-^ and PD-1^+^Tim-3^+^ CAR T-cells that received vaccine boosting (**Figure S4**). To assess the role of these cytokines in AS, we treated tumor-bearing mice with CAR T-vax therapy in the presence of neutralizing antibodies against IFN-γ or TNF-α. Therapy in the presence of isotype control or TNF-α-blocking antibodies had no impact on endogenous T-cell priming, but IFN-γ blockade completely abrogated priming of IFN-γ-producing endogenous anti-tumor T-cells (**Figure 5B**). To ensure that this effect did not reflect any potential artifact of the neutralizing antibodies interfering with *ex vivo* detection of cytokine-producing T-cells, and to rule out the possibility that tumor-specific T-cells were still being primed but had a different effector profile, we repeated IFN-γ blockade experiments in a second model of EGFRvIII^+^ CT-2A cells expressing OVA. CAR T-vax treatment expanded OVA-specific T-cells and also induced IFN-γ-producing T-cells recognizing SIINFEKL, as determined by peptide-MHC tetramer staining and ELISPOT, respectively (**Figure 5C-D**). However, IFN-γ neutralization during treatment eliminated the OVA-specific T-cell response by both readouts (**Figure 5C-D**). Further, endogenous T-cell infiltration and functional enhancement were also repressed by IFN-γ blockade during CAR T-vax treatment (**Figure S5, Supplemental Table 5**). Treatment of tumor-bearing animals with IFN-γ-deficient CAR T-cells also failed to elicit endogenous T-cell priming (**Figure 5E**), and knocking out PGC-1α significantly compromised IFN-γ production by intratumoral CAR T-cells (**Figure 5F**). These observations indicate that CAR T-cells are the major source of IFN-γ responsible for driving antigen spreading, sustained at least in part by OXPHOS. Administration of IFN-γ-neutralizing antibodies at different time points during therapy revealed that this cytokine was most critical for promoting AS during the first week of treatment (**Figure 5G**). While CAR T-vax therapy was highly effective in regressing CT-2A tumors, blockade of IFN-γ using neutralizing antibodies or treatment with IFN-γ-deficient CAR T-cells greatly reduced the efficacy of the treatment (**Figure 5H-J**). These data collectively point to a critical role for CAR T-cell-derived IFN-γ in promoting antigen spreading, which is amplified by vaccine boosting.

**Figure. 5.**
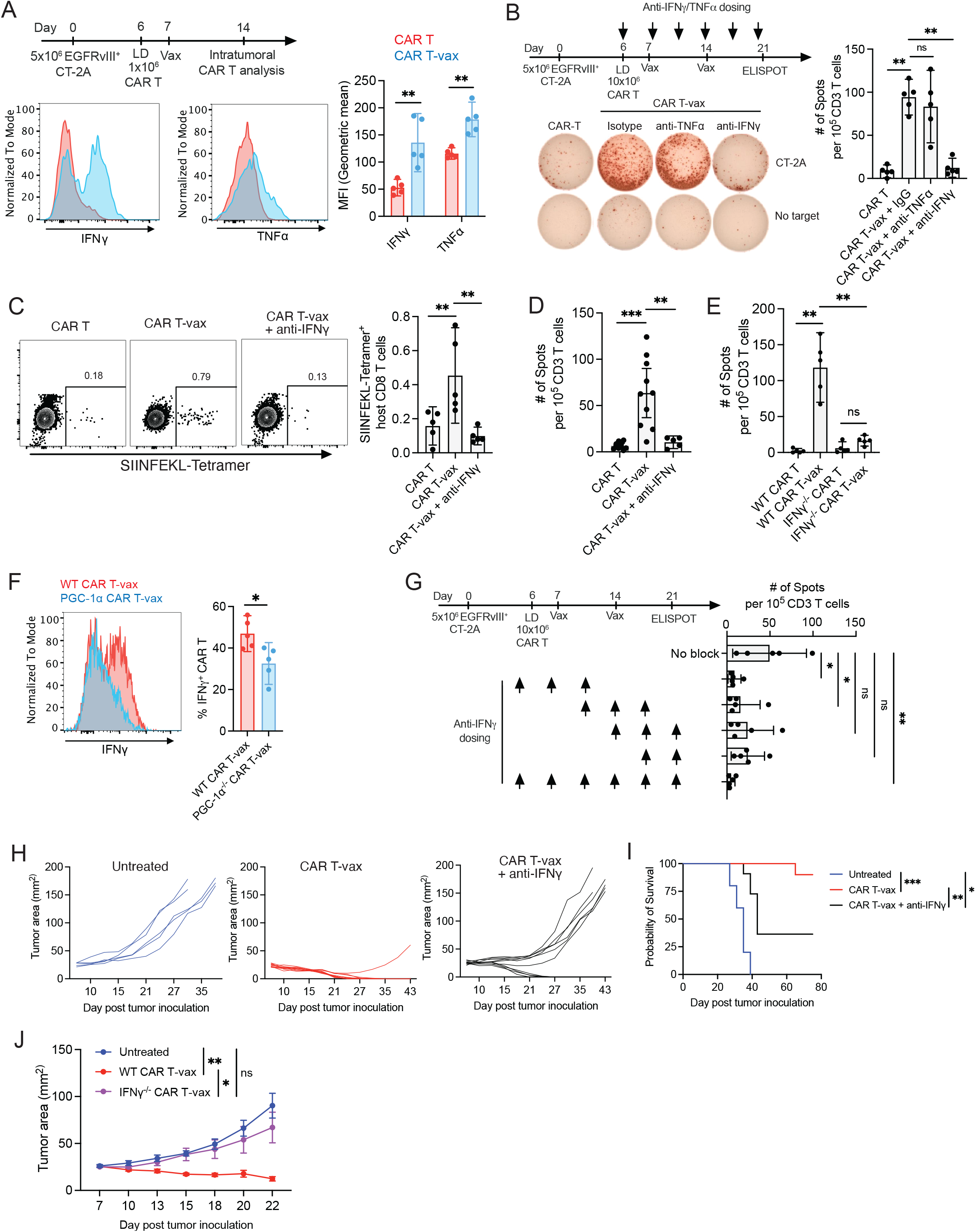
Enhanced IFN-γ production by vaccine-boosted CAR T-cells is critical for antigen spreading. (A) C57BL/6 mice bearing EGFRvIII^+^CT-2A tumors (*n* = 5 animals/group) were treated by EGFRvIII-CAR T ± vax and intratumoral CAR T-cells were analyzed for IFN-γ and TNF-α levels by flow cytometry on day 14 as shown in the timeline. Shown are representative flow histograms and geom. mean fluorescence intensities of cytokine staining. **, p < 0.01 by Student’s *t*-test. (B) CT-2A tumor-bearing mice (*n* = 5 animals/group) were treated with CAR-T or CAR T-vax in the presence of isotype control antibody (IgG) or neutralizing antibodies against TNF-α or IFN-γ. Antigen spreading was assessed by IFN-γ ELISPOT at day 21 on splenic T cells restimulated with irradiated EGFRvIII-CT-2A cells. Shown are representative ELISPOT well images and mean spots per sample. Experimental timeline and dosing intervals shown at top. Shown is one representative of at least three independent experiments. (C-D) C57BL/6 mice bearing OVA^+^EgFRvIII^+^CT-2A tumors were treated by EGFRvIII-CAR T or CAR T-vax ± neutralizing IFN-γ antibodies. Shown are endogenous OVA-specific T-cell responses on day 14 detected by flow cytometry analysis of SIINFEKL-tetramer staining (C) and ELISPOT on splenic T cells restimulated with irradiated OVA^+^ CT-2A cells (D). (E) CT-2A tumor-bearing mice (n = 5 animals/group) were treated with WT or IFN-γ^-/-^ CAR T ± vaccine boosting and endogenous T cell responses were analyzed by ELISPOT on day 21 as in panel (B). (F) CT-2A tumor-bearing mice (*n* = 5 animals/group) were treated with WT or PGC-1α^-/-^ CAR-T cells + vaccine boosting as in Fig. 5A, and IFN-γ expression was assessed in intratumoral CAR T-cells 7 days post CAR T-vax treatment by flow cytometry. *, p < 0.05 by Student’s *t*-test. Shown is one representative of two independent experiments. (G) CT-2A tumor-bearing mice (*n* = 5 animals/group) were treated with CAR T-vax and simultaneously received treatment with IFN-γ-neutralizing antibodies on days indicated by arrows; endogenous T cell responses in spleen were assessed by ELISPOT on day 21 as in B. (H-I) CT-2A tumor-bearing mice (*n* = 5 animals for untreated and 10 for CAR T-vax ± anti-IFNγ group) were treated with CAR-T vax ± IFN-γ-neutralizing antibodies or left untreated. Shown are individual tumor growth curves (H) and overall survival (I). ***, p<0.0001; **, p<0.01; *,p<0.05,; ns, not significant by Log-rank (Mantel-Cox) test. Shown is the combined data from two independent experiments. (J) CT-2A tumor-bearing mice (*n* = 5 animals for untreated and 10 for CAR T-vax ± anti-IFNγ group) were treated with WT or IFN^-/-^ CAR-T vax as in B. Shown is mean tumor growth. ns, not significant; *, p < 0.05; **, p < 0.01 by two-way ANOVA with Tukey’s post test. Shown is the combined data from two independent experiments. In panels B-D, and G, data shown is mean ± 95% CI, ***, p<0.0001; **p<0.01; *, p<0.05, ns, not significant by one-way ANOVA with Tukey’s post-test.

### IFN-γ sustains vaccine-boosted CAR T effector functions, promotes DC recruitment and antigen uptake, and triggers IL-12-mediated CAR-T-DC crosstalk

We next sought to define how CAR T-derived IFN-γ promoted antigen spreading, focusing on key steps in the chain of events leading to T-cell priming. Autocrine signaling from IFN-γ has been found to support the cytotoxicity of conventional T-cells (Bhat et al., 2017). To test if IFN-γ also promotes CAR-T killing in a similar manner, we evaluated the cytotoxicity of IFN-γ^-/-^ and IFNGR1^-/-^ CAR T-cells against EGFRvIII^+^ CT-2A cells *in vitro* and found that lack of IFN-γ or IFNGR1 expression by the CAR T-cells reduced cytotoxicity by ~50% (**Figure 6A**). Consistent with this finding, vaccine-boosted CAR-T that have elevated IFN-γ expression also exhibited increased granzyme B levels in tumors (**Figure S6A**) and tumor cells exhibited increased signatures of immunogenic cell death, such as upregulated cell surface calreticulin expression (**Figure S6B**).

**Figure. 6.**
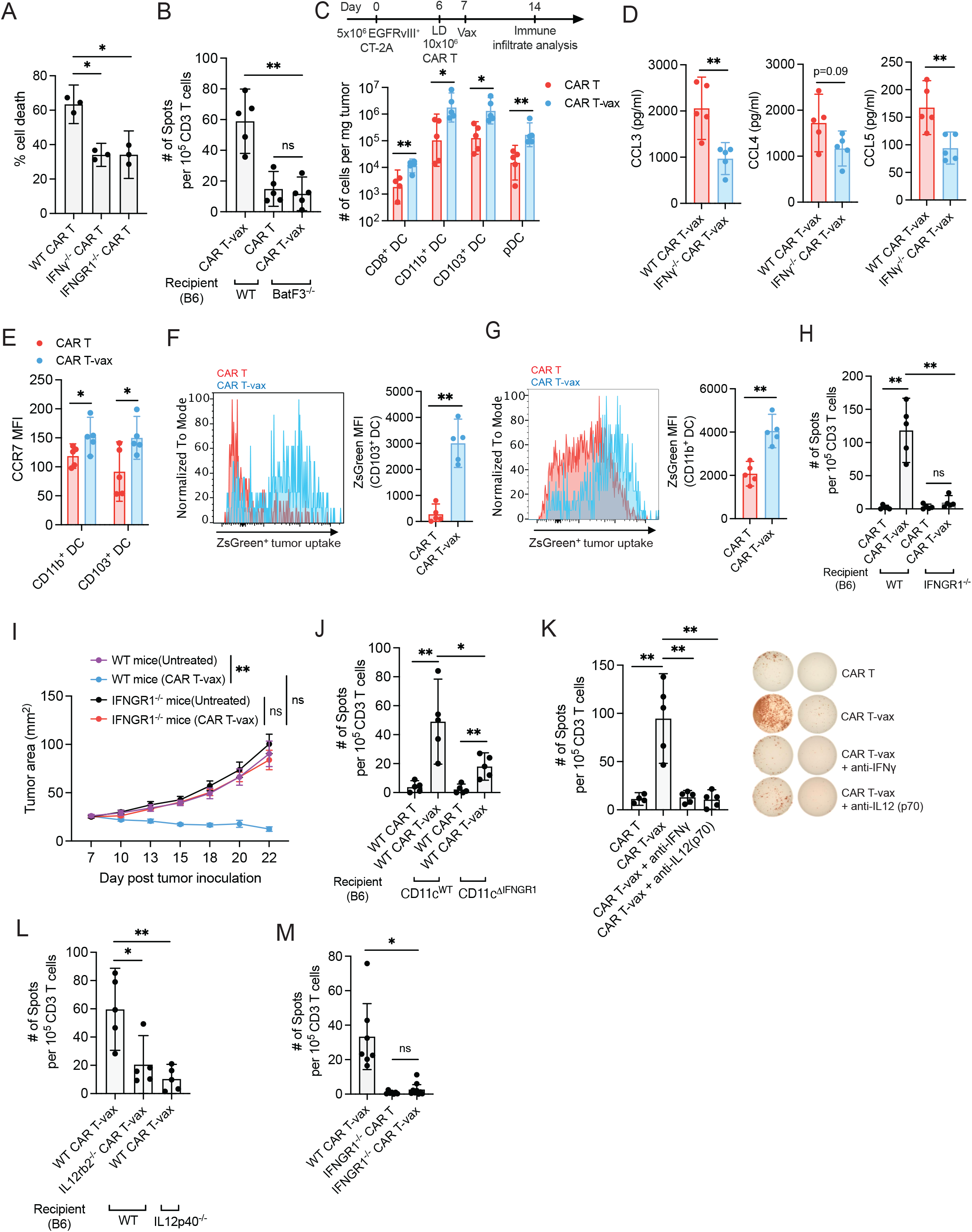
DCs regulate CAR T cell-induced antigen spreading through cross presentation and IFN-γ-IL-12 crosstalk. (A) EGFRvIII^+^ CT-2A cells were co-cultured with WT, IFN-γ^-/-^, or IFNGR1^-/-^ EGFRvIII-CAR T-cells *in vitro* at an 10:1 E:T ratio (*n* = 3 biol. replicates/group) and tumor cell death was measured by Sytox Red staining after 6 hr. Shown is one representative of at least three independent experiments. (B) WT or Batf3^-/-^ mice bearing CT-2A tumors were treated with CAR-T alone or CAR T-vax and endogenous T cell responses were assessed by IFN-γ ELISPOT on day 21 as in Fig. 1C. (C) CT-2A tumor-bearing mice were treated with CAR-T ± vaccine boosting. On day 14, intratumoral CD8^+^DCs, CD11b^+^DCs, CD103^+^DCs and plasmacytoid DCs (pDC) were enumerated. Shown is the combined data from two independent experiments. See supplemental methods for phenotyping details. (D) CT-2A tumor-bearing mice were treated with WT or IFN-γ^-/-^ CAR T cells + vaccine boosting. Chemokine expression in tumors 7 days post vaccine boosting was assessed by Luminex assay. (E) CT-2A tumor-bearing mice were treated with CAR-T ± vaccine boosting as in (C). CCR7 expression on intratumoral CD103^+^ DCs and CD11b^+^ DCs 7 days post T cell transfer was assessed by flow cytometry. (F-G) C57BL/6 mice bearing ZsGreen^+^EGFRvIII^+^ CT-2A tumors were treated as in (C) and intratumoral CD103^+^ DCs (G) and CD11b^+^ DCs (G) were analyzed on day 14 for tumor antigen (ZsGreen) uptake by flow cytometry. (H-I) CT-2A tumor-bearing mice were treated with WT or IFNGRV^-^ CAR T-cells ± vaccine boosting as in Fig. 1C. Splenic T cells were isolated on day 21 and restimulated with irradiated EGFRvIII-CT-2A cells for IFN-γ ELISPOT (H) and tumor growth was followed over time (I). **, p<0.01; ns, not significant. by two-way ANOVA with Tukey’s post test. (J) ELISPOT assay of endogenous T-cell responses as in (H) following CAR T-vax therapy in WT vs. CD11c-specific IFNGR1 KO mice. (K) ELISPOT assay of endogenous T-cell responses as in (H) following CAR T-vax therapy in the presence of neutralizing antibodies to IFN-γ or IL12(p70). (L) ELISPOT assay of endogenous T-cell responses as in (H) following WT or IL12rb2^-/-^ CAR T-vax therapy or in IL12p40^-/-^ mice receiving WT CAR T-vax therapy. (M) ELISPOT assay of endogenous T-cell responses as in (H) following WT or IFNGR1^-/-^ CAR T-vax therapy in WT tumor-bearing mice. Shown is one representative of two independent experiments. Throughout, n=5-6 animals/group. Data shown are mean ± 95% CI. **, p<0.01; *, p<0.05; ns, not significant by Student’s *t*-test (C-G) or one-way ANOVA followed by Tukey’s post test (A-B, H, J-M).

We next examined the DC compartment of treated tumors, as tumor antigen released by CAR T-mediated tumor killing must be acquired by DCs to drive endogenous T-cell priming. As expected, AS induced by CAR T-vax treatment was greatly reduced in *Batf3^-/-^* animals lacking cross-presenting DCs that are critical for anti-tumor T-cell priming (Böttcher and Sousa, 2018; Murphy and Murphy, 2022) (**Figure 6B**). Vaccine boosting of CAR T-cells led to substantial increases in multiple DC populations infiltrating treated tumors, including plasmacytoid DCs (pDCs), CD8^+^ DCs, CD103^+^ cDC1s (10-fold increase), and CD11b^+^ cDC2s (11-fold increase) (**Figure 6C**). DC recruitment to tumor relies on chemokines such as CCL3, CCL4 and CCL5 (Böttcher et al., 2018; Vilgelm and Richmond, 2019), and intratumoral expression of these chemokines was reduced when treating with IFN-γ^-/-^ CAR T-cells (**Figure 6D**). Using an EGFRvIII^+^CT-2A tumor line expressing ZsGreen as a traceable antigen, we found that vaccine boosting triggered upregulation of the lymph node homing marker CCR7 (**Figure 6E, S6C-D**) and increased tumor antigen uptake by both cDC1 and cDC2 populations in the tumor, notably by ~11-fold for cDC1s (**Figure 6F-G**). Thus vaccine-boosting CAR T-cells amplified multiple prerequisite steps for antigen spreading.

Our *in vitro* analysis suggested a role for autocrine CAR T-cell-derived IFN-γ in sustaining CAR T cytotoxicity, but the target cells responding to IFN-γ *in vivo* remained unclear. We first tested if host cells were important responders, by transferring CAR T-cells into tumor-bearing WT or IFNGR1’^/_^ mice, followed by vaccine boosting. Endogenous T-cell priming and tumor control were nearly completely lost in IFNGR1’^/_^ mice (**Figure 6H-I**). Next, we generated mice with specific deletion of IFNGR1 in CD11c^+^ DCs by crossing CD11c-cre and IFNGR-floxed animals to generate CD11c^ΔIFNGR1^ mice. As shown in **Figure 6J**, CAR T-vax treatment of tumor-bearing CD11c^ΔIFNGR1^ mice led to reduced by not fully ablated endogenous T-cell priming, suggesting that DCs are important responders but not the sole host cell population stimulated by IFN-γ. Activation of dendritic cells by T-cell-derived IFN-γ has been shown to trigger production of IL-12 by DCs, which in turn acts as positive feedback signal reinforcing T-cell IFN-γ expression and cytotoxic activity during checkpoint blockade immunotherapy (Garris et al., 2018). Strikingly, antibody-mediated neutralization of IL-12 during CAR T-vax therapy eliminated antigen spreading comparably to IFN-γ blockade (**Figure 6K**). A similar loss of AS was observed when CAR T-vax therapy was administered to IL-12-deficient mice (**Figure 6L**). The CAR T-cells themselves are important responders to IL-12 as therapy with IL-12Rb2^-/-^ CAR T-cells elicited nearly baseline endogenous T-cell priming in 4 of 5 animals (**Figure 6L**).

IL-12 drives sustained/elevated autocrine IFN-γ expression by T-cells. *In vivo,* vaccine-boosted IFNGR1-deficient CAR T-cells showed reduced production of IFN-γ, Granzyme B and a trend toward reduced levels of TNF-α (**Figure S7A-D**). Blunted effector functions of IFNGR1-deficient CAR T-cells correlated with reduced induction of immunogenic cell death markers on tumor cells (**Figure S7E**), decreased tumor antigen uptake by intratumoral cDC1 and cDC2s (**Figure S7F-G**), and reduced tumor antigen acquisition by lymph node-resident CD8α^+^ cDC1 (**Figure S7H**); tumor antigen uptake by LN cDC2 was low and unaffected (**Figure S7I**). These changes in CAR-T function, tumor cell killing and tumor antigen release correlated with complete loss of endogenous T-cell priming when tumor-bearing animals were treated with CAR T-vax therapy using IFNGR1^-/-^ CAR T-cells (**Figure 6M**). Altogether, vaccine boosting enables CAR T-cells to sustain cytotoxicity in the TME and drive key events required for antigen spreading, dependent both on the ability of host DCs and the CAR T-cells themselves to respond IFN-γ.

### Robust IFN-γ production is essential for CAR T-vax therapy to control tumors with pre-existing antigen heterogeneity

Based on our collected mechanistic findings regarding the importance of IFN-γ in driving antigen spreading and supporting effector functions of both CAR and endogenous T-cells, we finally assessed the importance of sustained cytokine production by CAR T-cells in promoting control of antigenically heterogeneous tumors. We focused on the treatment case of tumors generated from mixtures of 80% EGFRvIII^+^ and 20% EGFRvIII^-^ tumor cells. Unsurprisingly, CAR T-vax therapy in the presence of IFN-γ blockade led to loss of survival extension and elicited no complete responses (**Figure 7A)**. We hypothesized that enforced expression of IFN-γ might further enhance endogenous T-cell priming elicited by CAR T-vax therapy. To test this idea, we transduced CAR T-cells with retroviral constructs bearing an NFAT-driven IFN-γ expression cassette, to obtain elevated IFN-γ production following CAR activation (Zhang et al., 2011). We confirmed that NFAT-IFN-γ CAR T-cells produced nearly twice as much of IFN-γ as WT CAR T-cells upon stimulation *in vitro* (**Figure 7B**). Non-vaccine boosted NFAT-IFN-γ CAR T therapy itself elicited a significant level of endogenous T-cell priming, consistent with a critical role for sustained CAR T-cell IFN-γ in AS generally (**Figure 7C**). AS was further increased when NFAT-IFN-γ CART were used in combination with vaccine boosting, reaching 50% higher levels than treatment with WT CAR T-cells (**Figure 7C**). We then assessed whether enforced IFN-γ expression could enhance control of heterogeneous tumors. Vaccine-boosted WT CAR T-cells were able to reject 25-50% of 80:20 EGFRvIII^+^:EGFRvIII^-^ mixed tumors (**Figure 7A, D-E**). NFAT-IFN-γ CAR T-cells achieved similar complete response rates in the 20% antigen-negative tumor model in the absence of vaccine boosting, and strikingly, this complete response rate increased to 80% when vaccine boosting was added to the treatment (**Figure 7D-E**). Thus, strategies to enhance IFN-γ production and favorable CAR T-cell metabolism appear promising to increase the efficacy of CAR T-cell therapy against antigenically heterogenous solid tumors.

**Figure. 7.**
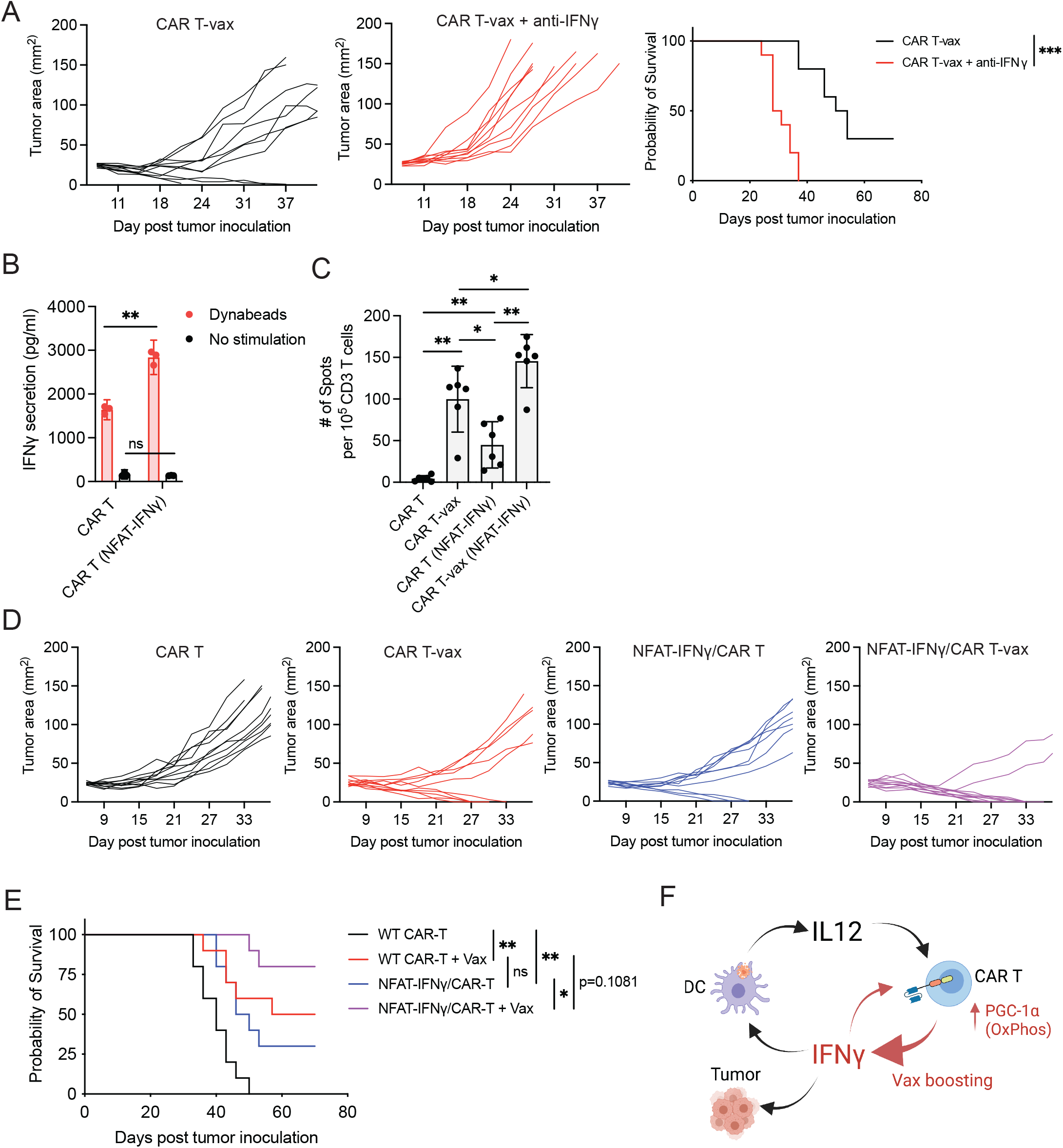
Engineering CAR T-cells for increased IFN-γ expression synergizes with vaccine boosting to enhance antigen spreading and rejection of solid tumors with pre-existing antigen heterogeneity. (A) Mice bearing heterogenous tumors were established by inoculating a mixture of EGFRvIII^+^ CT-2A and WT CT-2A cells at an 80:20 ratio into the flank of C57BL/6 mice. Shown are tumor growth and survival of animals receiving WT CAR T-vax therapy in the presence or absence of IFN-γ antibodies. Shown is the combined data from two independent experiments. (B) EGFRvIII-CAR T-cells or CAR-T transduced with an NFAT-driven mouse IFN-γ expression cassette were cultured with or without anti-CD3/CD28 beads for 6 hr, and IFN-γ released in the supernatant was measured by ELISA (*n*=3 biol. replicates/group). (C) ELISPOT assay as in Fig. 6H showing endogenous T-cell responses induced in CT-2A tumor-bearing mice following treatment with WT or NFAT-IFN-γ-transduced CAR T ± vaccine boosting. (D-E) CT-2A tumor-bearing mice were treated as in Fig. 1C with WT or NFAT-IFN-γ-transduced CAR T-cells in the presence or absence of vaccine boosting. Shown are Tumor growth (D) and overall survival (E). ***, p<0.0001; **, p<0.01; *, p<0.05 by two-way ANOVA with Tukey’s post test (tumor growth) or Log-rank (Mantel-Cox) test (overall survival). Shown is the combined data from two independent experiments. (F) Schematic view of vaccine-triggered metabolic reprogramming of CAR T-cells and IFN-γ-IL12 crosstalk between CAR T and DCs in tumors. In panels A, D-E, *n*=10 animals/group. In panel C, *n*=6 animals/group. In panel B-C, data shown is mean ± 95% CI. **, p<0.01; *, p<0.05; ns not significant by one-way ANOVA followed by Tukey’s post test (C).

## Discussion

Antigenic heterogeneity and antigen loss play important roles in tumor escape from immune surveillance and represent mechanisms of resistance to antigen-specific CAR-T therapies (Guedan et al., 2019; Lim and June, 2017; Majzner and Mackall, 2018). The induction of antigen spreading by CAR-T therapy would be an attractive approach to overcome these challenges. However, evidence for AS in response to CAR T treatment alone in humans remains limited. Further, preclinical studies using combination therapies or CAR T-cells transduced with one or more supporting genes have reported induction of AS and endogenous anti-tumor T-cell responses, but mechanisms governing these responses remain poorly understood. Here we demonstrate that CAR T-cells bearing second-generation CARs, which receive *in vivo* restimulation via a vaccine activating the CAR in lymph nodes, are capable of promoting robust antigen spreading with induction of both CD4^+^ and CD8^+^ T-cell responses against non-CAR-related tumor antigens. This endogenous T-cell response has significant consequences for the outcome of CAR T therapy: (1) long-term tumor regressions and complete responses are achieved against tumors that otherwise undergo antigen loss-based relapse; (2) control of antigenically heterogeneous tumors can be achieved, with a few animals rejecting even tumors that are 50% comprised of CAR antigen-negative cancer cells at the time of treatment; (3) long term protection against tumor rechallenge is achieved.

To understand how vaccine boosting enabled CAR T-driven antigen spreading, we examined changes in CAR T functionality, transcriptional profiles, and alterations in the TME induced in response to CAR T-vax therapy. These studies revealed enhanced production of IFN-γ by vaccine-boosted CAR T-cells as a major contributor to the endogenous T-cell priming. In natural immune responses, IFN-γ is a critical effector cytokine produced by T-cells promoting the activation of both innate and adaptive immunity (Schroder et al., 2004), maintenance of T-cell cytotoxicity and mobility (Bhat et al., 2017), polarization of T helper cells to Th1 cells (Castro et al., 2018), reduction of Treg-mediated suppression (Overacre-Delgoffe et al., 2017) and sensitization of tumor cells to T-cell-mediated cytotoxicity (Castro et al., 2018). However, the role of IFN-γ in the function of CAR T-cells remains poorly defined. Recently, IFN-γ has been reported to preferentially regulate the expression of cell adhesion molecules on solid tumor cells, but not leukemic cells, and subsequently enhance CAR T-cell-mediated cytotoxicity by stabilizing CAR T-tumor cell engagement (Larson et al., 2022). Here we show that IFN-γ production by CAR T-cells and sensing of IFN-γ by both host cells and the CAR T-cells themselves are also critical to enable an antigen spreading response. Mechanistically, IFN-γ played an important role in maintaining sustained high levels of cytotoxicity and effector cytokine expression in vaccine-boosted CAR T-cells in a cell-intrinsic manner. These enhanced CAR T-cell effector functions in turn correlated with increased expression of DC-recruiting chemokines in tumors, increased DC infiltration, tumor antigen uptake, and activation of intratumoral DCs. These effects of CAR T-derived IFN-γ were propagated via a positive feedback regulation through crosstalk with DC-derived IL-12. The observation that defective IL-12 signaling in both CAR-T-cells and the host phenocopies IFN-γ deficiency in abrogating AS suggests that CAR T-derived IFN-γ acts predominantly as a trigger for prolonged IFN-γ production, driven by DC-derived IL-12 in the TME. Such IFN-γ/IL-12 crosstalk has proven to underlie a number of successful immunotherapies, including checkpoint blockade therapy (Garris et al., 2018) and CAR T-cell therapy in lymphoma (Boulch et al., 2021). Our data do not exclude potential contributions of other important cytokines or immune cell types, such as tumor-resident macrophages, the activation of which has been reported to be an important mechanism how CAR T-cells engage endogenous immunity (Alizadeh et al., 2021).

IFN-γ production is tightly regulated at both the transcriptional level by transcription factors (TFs) (Samten et al., 2008; Xu et al., 2019) including CREB, AP-1, T-bet, NFAT, and at the post-transcriptional level by various miRNAs, ARE or GAPDH binding to its 3’UTR (Chang et al., 2013; Savan, 2014). Although IFN-γ synthesis has been proposed to be predominantly associated with glycolysis due to its regulation by GAPDH (Chang et al., 2013), both glycolysis and oxidative phosphorylation were proven crucial in controlling IFN-γ production in NK cells (Donnelly et al., 2014; Keating et al., 2016), consistent with previous reports that elevated OXPHOS and mitochondria integrity was required to support IFN-γ production (Gerbec et al., 2020; Keating et al., 2016; Lisci et al., 2021). Thus, it seems likely that vaccination-induced metabolic reprogramming of CAR T-cells including increased OXPHOS support the sustained IFN-γ production achieved by vaccine-boosted CAR T-cells in the TME. This is in line with our observation that genetic deletion of PGC-1α, a key TF regulating OXPHOS, resulted in reduced expression of IFN-γ and a significant reduction in AS. OXPHOS is often an important feature of memory-like T-cells (Windt and Pearce, 2012), and enforced expression of PGC-1α endows T-cells with superior anti-tumor activity (Scharping et al., 2016). Despite the extensive elevation of multiple metabolic pathways in response to vaccine boosting, including OXPHOS, mTORC1, Fatty-acid metabolism, and peroxisome signaling, we cannot rule out the possibility that multiple metabolic pathways (including glycolysis) collectively contribute to sustained IFN-γ production in intratumoral CAR T-cells. The extent to which each metabolic pathway and which gene(s), including GAPDH, are responsible for IFN-γ production by vaccine-boosted intratumoral CAR T-cells will require future work.

In summary, we have shown that vaccine boosting through the chimeric receptor triggers markedly enhanced CAR T polyfunctionality and metabolic reprogramming (**Figure 7F**). Vaccine-boosted CAR T-cells trigger robust recruitment and activation of DCs in the tumor, which in turn secrete IL-12 that, together with the autocrine effect of IFN-γ, enhances CAR T-cell anti-tumor activity, leading to enhanced AS. Few solid tumors express target antigens on >90% of tumor cells. The findings described here provide guidance for engineering CAR-T therapies to more effectively treat solid tumors with pre-existing antigenic heterogeneity.

## ACKNOWLEDGMENTS

We thank the Koch Institute Swanson Biotechnology Center for technical support, specifically the flow cytometry core facility. We thank Dr. Thomas Seyfried for providing CT-2A cell line.

## Data and materials availability

All data are available on request.

## AUTHOR CONTRIBUTIONS

L.M., D.J.I. and J.C.L designed the studies. L.M. D.M.M, D.J.I. analyzed and interpreted the data and wrote the manuscript. L.M. performed the experiments. L.M. carried out amphiphile-peptide vaccine synthesis. I.S. assisted with CAR T production, sample preparation, and ELISPOT assays. D.M.M, and C.W. perform single cell and bulk RNA-sequencing analysis. P.Y assisted with Env/p15E antigen validation. W.A., and N.L. assisted in assay preparation.

## REFERENCES

Adachi, K., Kano, Y., Nagai, T., Okuyama, N., Sakoda, Y., and Tamada, K. (2018). IL-7 and CCL19 expression in CAR-T cells improves immune cell infiltration and CAR-T cell survival in the tumor. Nature Biotechnology 36, 346–351.

Alizadeh, D., Wong, R.A., Gholamin, S., Maker, M., Aftabizadeh, M., Yang, X., Pecoraro, J.R., Jeppson, J.D., Wang, D., Aguilar, B., et al. (2021). IFNγ is Critical for CAR T Cell–mediated Myeloid Activation and Induction of Endogenous Immunity. Cancer Discov 11, 2248–2265.

Awad, M.M., Govindan, R., Balogh, K.N., Spigel, D.R., Garon, E.B., Bushway, M.E., Poran, A., Sheen, J.H., Kohler, V., Esaulova, E., et al. (2022). Personalized neoantigen vaccine NEO-PV-01 with chemotherapy and anti-PD-1 as first-line treatment for non-squamous non-small cell lung cancer. Cancer Cell 40, 1010–1026.e11.

Beatty, G.L., Haas, A.R., Maus, M.V., Torigian, D.A., Soulen, M.C., Plesa, G., Chew, A., Zhao, Y., Levine, B.L., Albelda, S.M., et al. (2014). Mesothelin-Specific Chimeric Antigen Receptor mRNA-Engineered T Cells Induce Antitumor Activity in Solid Malignancies. Cancer Immunol Res 2, 112–120.

Bhat, P., Leggatt, G., Waterhouse, N., and Frazer, I.H. (2017). Interferon-γ derived from cytotoxic lymphocytes directly enhances their motility and cytotoxicity. Cell Death Dis 8, e2836–e2836.

Böttcher, J.P., and Sousa, C.R. e (2018). The Role of Type 1 Conventional Dendritic Cells in Cancer Immunity. Trends Cancer 4, 784–792.

Böttcher, J.P., Bonavita, E., Chakravarty, P., Blees, H., Cabeza-Cabrerizo, M., Sammicheli, S., Rogers, N.C., Sahai, E., Zelenay, S., and Sousa, C.R. e (2018). NK Cells Stimulate Recruitment of cDC1 into the Tumor Microenvironment Promoting Cancer Immune Control. Cell 172, 1022–1037.e14.

Boulch, M., Cazaux, M., Loe-Mie, Y., Thibaut, R., Corre, B., Lemaître, F., Grandjean, C.L., Garcia, Z., and Bousso, P. (2021). A cross-talk between CAR T cell subsets and the tumor microenvironment is essential for sustained cytotoxic activity. Sci Immunol 6.

Brossart, P. (2020). The role of antigen-spreading in the efficacy of immunotherapies. Clin Cancer Res clincanres.0305.2020.

Castro, F., Cardoso, A.P., Gonçalves, R.M., Serre, K., and Oliveira, M.J. (2018). Interferon-Gamma at the Crossroads of Tumor Immune Surveillance or Evasion. Front Immunol 9, 847.

Chang, C.-H., Curtis, J.D., Maggi, L.B., Faubert, B., Villarino, A.V., O’Sullivan, D., Huang, S.C.-C., van der Windt, G.J.W., Blagih, J., Qiu, J., et al. (2013). Posttranscriptional Control of T Cell Effector Function by Aerobic Glycolysis. Cell 153, 1239–1251.

Chmielewski, M., and Abken, H. (2017). CAR T Cells Releasing IL-18 Convert to T-Bethigh FoxO1low Effectors that Exhibit Augmented Activity against Advanced Solid Tumors. Cell Reports 21, 3205–3219.

Donnelly, R.P., Loftus, R.M., Keating, S.E., Liou, K.T., Biron, C.A., Gardiner, C.M., and Finlay, D.K. (2014). mTORC1-Dependent Metabolic Reprogramming Is a Prerequisite for NK Cell Effector Function. J Immunol 193, 4477–4484.

Etxeberria, I., Bolaños, E., Quetglas, J.I., Gros, A., Villanueva, A., Palomero, J., Sánchez-Paulete, A.R., Piulats, J.M., Matias-Guiu, X., Olivera, I., et al. (2019). Intratumor Adoptive Transfer of IL-12 mRNA Transiently Engineered Antitumor CD8+ T Cells. Cancer Cell 36, 613–629.e7.

Fernandez-Marcos, P.J., and Auwerx, J. (2011). Regulation of PGC-1α, a nodal regulator of mitochondrial biogenesis. Am J Clin Nutrition 93, 884S–890S.

Fesnak, A.D., June, C.H., and Levine, B.L. (2016). Engineered T cells: the promise and challenges of cancer immunotherapy. Nat Rev Cancer 16, 566–581.

Garris, C.S., Arlauckas, S.P., Kohler, R.H., Trefny, M.P., Garren, S., Piot, C., Engblom, C., Pfirschke, C., Siwicki, M., Gungabeesoon, J., et al. (2018). Successful Anti-PD-1 Cancer Immunotherapy Requires T Cell-Dendritic Cell Crosstalk Involving the Cytokines IFN-γ and IL-12. Immunity 49, 1148–1161.e7.

Gerbec, Z.J., Hashemi, E., Nanbakhsh, A., Holzhauer, S., Yang, C., Mei, A., Tsaih, S.-W., Lemke, A., Flister, M.J., Riese, M.J., et al. (2020). Conditional Deletion of PGC-1α Results in Energetic and Functional Defects in NK Cells. Iscience 23, 101454.

Guedan, S., Calderon, H., Jr., A.D.P., and Maus, M.V. (2019). Engineering and Design of Chimeric Antigen Receptors. Molecular Therapy - Methods & Clinical Development 12, 145–156.

Gulley, J.L., Madan, R.A., Pachynski, R., Mulders, P., Sheikh, N.A., Trager, J., and Drake, C.G. (2017). Role of Antigen Spread and Distinctive Characteristics of Immunotherapy in Cancer Treatment. Jnci J National Cancer Inst 109, djw261.

Hegde, M., Joseph, S.K., Pashankar, F., DeRenzo, C., Sanber, K., Navai, S., Byrd, T.T., Hicks, J., Xu, M.L., Gerken, C., et al. (2020). Tumor response and endogenous immune reactivity after administration of HER2 CAR T cells in a child with metastatic rhabdomyosarcoma. Nat Commun 11, 3549.

Hou, A.J., Chen, L.C., and Chen, Y.Y. (2021). Navigating CAR-T cells through the solid-tumour microenvironment. Nat Rev Drug Discov 20, 531–550.

Keating, S.E., Zaiatz-Bittencourt, V., Loftus, R.M., Keane, C., Brennan, K., Finlay, D.K., and Gardiner, C.M. (2016). Metabolic Reprogramming Supports IFN-γ Production by CD56bright NK Cells. J Immunol 196, 2552–2560.

Kerkar, S.P., Muranski, P., Kaiser, A., Boni, A., Sanchez-Perez, L., Yu, Z., Palmer, D.C., Reger, R.N., Borman, Z.A., Zhang, L., et al. (2010). Tumor-Specific CD8+ T Cells Expressing Interleukin-12 Eradicate Established Cancers in Lymphodepleted Hosts. Cancer Res 70, 6725–6734.

Khalil, D.N., Smith, E.L., Brentjens, R.J., and Wolchok, J.D. (2016). The future of cancer treatment: immunomodulation, CARs and combination immunotherapy. Nat Rev Clin Oncol 13, 273–290.

Kim, R.H., Plesa, G., Gladney, W., Kulikovskaya, I., Levine, B.L., Lacey, S.F., June, C.H., Melenhorst, J.J., and Beatty, G.L. (2017). Effect of chimeric antigen receptor (CAR) T cells on clonal expansion of endogenous non-CAR T cells in patients (pts) with advanced solid cancer. J Clin Oncol 35, 3011–3011.

Klampatsa, A., Leibowitz, M.S., Sun, J., Liousia, M., Arguiri, E., and Albelda, S.M. (2020). Analysis and Augmentation of the Immunologic Bystander Effects of CAR T Cell Therapy in a Syngeneic Mouse Cancer Model. Mol Ther Oncolytics 18, 360–371.

Kueberuwa, G., Kalaitsidou, M., Cheadle, E., Hawkins, R.E., and Gilham, D.E. (2018). CD19 CAR T Cells Expressing IL-12 Eradicate Lymphoma in Fully Lymphoreplete Mice through Induction of Host Immunity. Molecular Therapy: Oncolytics 8, 41–51.

Kuhn, N.F., Lopez, A.V., Li, X., Cai, W., Daniyan, A.F., and Brentjens, R.J. (2020). CD103+ cDC1 and endogenous CD8+ T cells are necessary for improved CD40L-overexpressing CAR T cell antitumor function. Nat Commun 11, 6171.

Kvistborg, P., Philips, D., Kelderman, S., Hageman, L., Ottensmeier, C., Joseph-Pietras, D., Welters, M.J.P., Burg, S. van der, Kapiteijn, E., Michielin, O., et al. (2014). Anti–CTLA-4 therapy broadens the melanoma-reactive CD8+ T cell response. Sci Transl Med 6, 254ra128.

Lai, J., Mardiana, S., House, I.G., Sek, K., Henderson, M.A., Giuffrida, L., Chen, A.X.Y., Todd, K.L., Petley, E.V., Chan, J.D., et al. (2020). Adoptive cellular therapy with T cells expressing the dendritic cell growth factor Flt3L drives epitope spreading and antitumor immunity. Nat Immunol 1–13.

Landsberg, J., Kohlmeyer, J., Renn, M., Bald, T., Rogava, M., Cron, M., Fatho, M., Lennerz, V., Wölfel, T., Hölzel, M., et al. (2012). Melanomas resist T-cell therapy through inflammation-induced reversible dedifferentiation. Nature 490, 412–416.

Larson, R.C., Kann, M.C., Bailey, S.R., Haradhvala, N.J., Llopis, P.M., Bouffard, A.A., Scarfó, I., Leick, M.B., Grauwet, K., Berger, T.R., et al. (2022). CAR T cell killing requires the IFNγR pathway in solid but not liquid tumours. Nature 604, 563–570.

LeBleu, V.S., O’Connell, J.T., Herrera, K.N.G., Wikman, H., Pantel, K., Haigis, M.C., Carvalho, F.M. de, Damascena, A., Chinen, L.T.D., Rocha, R.M., et al. (2014). PGC-1α mediates mitochondrial biogenesis and oxidative phosphorylation in cancer cells to promote metastasis. Nat Cell Biol 16, 992–1003.

Lim, W.A., and June, C.H. (2017). The Principles of Engineering Immune Cells to Treat Cancer. Cell 168, 724–740.

Lisci, M., Barton, P.R., Randzavola, L.O., Ma, C.Y., Marchingo, J.M., Cantrell, D.A., Paupe, V., Prudent, J., Stinchcombe, J.C., and Griffiths, G.M. (2021). Mitochondrial translation is required for sustained killing by cytotoxic T cells. Science 374, eabe9977.

Liu, H., Moynihan, K.D., Zheng, Y., Szeto, G.L., Li, A.V., Huang, B., Egeren, D.S.V., Park, C., and Irvine, D.J. (2014). Structure-based programming of lymph-node targeting in molecular vaccines. Nature 507, 519–522.

Ma, L., Dichwalkar, T., Chang, J.Y.H., Cossette, B., Garafola, D., Zhang, A.Q., Fichter, M., Wang, C., Liang, S., Silva, M., et al. (2019). Enhanced CAR-T cell activity against solid tumors by vaccine boosting through the chimeric receptor. Science 365, 162–168.

Majzner, R.G., and Mackall, C.L. (2018). Tumor Antigen Escape from CAR T-cell Therapy. Cancer Discov 8, 1219–1226.

Murphy, T.L., and Murphy, K.M. (2022). Dendritic cells in cancer immunology. Cell Mol Immunol 19, 3–13.

O’Rourke, D.M., Nasrallah, M.P., Desai, A., Melenhorst, J.J., Mansfield, K., Morrissette, J.J.D., Martinez-Lage, M., Brem, S., Maloney, E., Shen, A., et al. (2017). A single dose of peripherally infused EGFRvIII-directed CAR T cells mediates antigen loss and induces adaptive resistance in patients with recurrent glioblastoma. Sci Transl Med 9, eaaa0984.

Overacre-Delgoffe, A.E., Chikina, M., Dadey, R.E., Yano, H., Brunazzi, E.A., Shayan, G., Horne, W., Moskovitz, J.M., Kolls, J.K., Sander, C., et al. (2017). Interferon-γ Drives Treg Fragility to Promote Anti-tumor Immunity. Cell 169, 1130–1141.e11.

Park, A.K., Fong, Y., Kim, S.-I., Yang, J., Murad, J.P., Lu, J., Jeang, B., Chang, W.-C., Chen, N.G., Thomas, S.H., et al. (2020). Effective combination immunotherapy using oncolytic viruses to deliver CAR targets to solid tumors. Sci Transl Med 12.

Rafiq, S., Hackett, C.S., and Brentjens, R.J. (2020). Engineering strategies to overcome the current roadblocks in CAR T cell therapy. Nat Rev Clin Oncol 17, 147–167.

Roselli, E., Faramand, R., and Davila, M.L. (2021). Insight into next-generation CAR therapeutics: designing CAR T cells to improve clinical outcomes. J Clin Invest 131, e142030.

Samten, B., Townsend, J.C., Weis, S.E., Bhoumik, A., Klucar, P., Shams, H., and Barnes, P.F. (2008). CREB, ATF, and AP-1 Transcription Factors Regulate IFN-γ Secretion by Human T Cells in Response to Mycobacterial Antigen. J Immunol 181, 2056–2064.

Savan, R. (2014). Post-Transcriptional Regulation of Interferons and Their Signaling Pathways. J Interf Cytokine Res 34, 318–329.

Scharping, N.E., Menk, A.V., Moreci, R.S., Whetstone, R.D., Dadey, R.E., Watkins, S.C., Ferris, R.L., and Delgoffe, G.M. (2016). The Tumor Microenvironment Represses T Cell Mitochondrial Biogenesis to Drive Intratumoral T Cell Metabolic Insufficiency and Dysfunction. Immunity 45, 374–388.

Schroder, K., Hertzog, P.J., Ravasi, T., and Hume, D.A. (2004). Interferon-γ: an overview of signals, mechanisms and functions. J Leukocyte Biol 75, 163–189.

Shah, N.N., and Fry, T.J. (2019). Mechanisms of resistance to CAR T cell therapy. Nature Reviews Clinical Oncology 1–14.

Szabo, P.A., Levitin, H.M., Miron, M., Snyder, M.E., Senda, T., Yuan, J., Cheng, Y.L., Bush, E.C., Dogra, P., Thapa, P., et al. (2019). Single-cell transcriptomics of human T cells reveals tissue and activation signatures in health and disease. Nat Commun 10, 4706.

Tibbitt, C.A., Stark, J.M., Martens, L., Ma, J., Mold, J.E., Deswarte, K., Oliynyk, G., Feng, X., Lambrecht, B.N., Bleser, P.D., et al. (2019). Single-Cell RNA Sequencing of the T Helper Cell Response to House Dust Mites Defines a Distinct Gene Expression Signature in Airway Th2 Cells. Immunity 51, 169–184.e5.

Vardhana, S.A., Hwee, M.A., Berisa, M., Wells, D.K., Yost, K.E., King, B., Smith, M., Herrera, P.S., Chang, H.Y., Satpathy, A.T., et al. (2020). Impaired mitochondrial oxidative phosphorylation limits the self-renewal of T cells exposed to persistent antigen. Nat Immunol 21, 1022–1033.

Vilgelm, A.E., and Richmond, A. (2019). Chemokines Modulate Immune Surveillance in Tumorigenesis, Metastasis, and Response to Immunotherapy. Front Immunol 10, 333.

Walsh, S.R., Simovic, B., Chen, L., Bastin, D., Nguyen, A., Stephenson, K., Mandur, T.S., Bramson, J.L., Lichty, B.D., and Wan, Y. (2019). Endogenous T cells prevent tumor immune escape following adoptive T cell therapy. J Clin Invest 129, 5400–5410.

Windt, G.J.W., and Pearce, E.L. (2012). Metabolic switching and fuel choice during T-cell differentiation and memory development. Immunol Rev 249, 27–42.

Xu, T., Keller, A., and Martinez, G.J. (2019). NFAT1 and NFAT2 Differentially Regulate CTL Differentiation Upon Acute Viral Infection. Front Immunol 10, 184.

Zhang, L., Kerkar, S.P., Yu, Z., Zheng, Z., Yang, S., Restifo, N.P., Rosenberg, S.A., and Morgan, R.A. (2011). Improving Adoptive T Cell Therapy by Targeting and Controlling IL-12 Expression to the Tumor Environment. Mol Ther 19, 751–759.

Zhang, L., Morgan, R.A., Beane, J.D., Zheng, Z., Dudley, M.E., Kassim, S.H., Nahvi, A.V., Ngo, L.T., Sherry, R.M., Phan, G.Q., et al. (2015). Tumor-Infiltrating Lymphocytes Genetically Engineered with an Inducible Gene Encoding Interleukin-12 for the Immunotherapy of Metastatic Melanoma. Clin Cancer Res 21, 2278–2288.

