## Supplemental materials for "Eradication of tumors with pre-existing antigenic heterogeneity by vaccine-mediated co-engagement of CAR T and endogenous T-cells"

**KEY RESOURCES TABLE**

| REAGENT or RESOURCE | SOURCE | IDENTIFIER |
|---|---|---|
| **Antibodies** | | |
| Anti-mouse CD3 (17A2) Alex488 | BioLegend | 100220; RRID:AB_1732057 |
| Anti-mouse CD8a (53-6.7) BUV395 | BD Biosciences | 563786; RRID:AB_2732919 |
| Anti-mouse CD8a (53-6.7) BV421 | BioLegend | 100738; RRID:AB_11204079 |
| Anti-mouse CD4 (RM4-5) FITC | BioLegend | 100510; RRID:AB_312713 |
| Anti-mouse CD25 (PC61) APC-Cy7 | BioLegend | 102026; RRID:AB_830745 |
| Anti-mouse B220 (RA3-6B2) PE-cy7 | BioLegend | 103222; RRID: AB_313005 |
| Anti-mouse PD-1 (29F.1A12) BV421 | BioLegend | 135218; RRID:AB_2561447 |
| Anti-mouse TIM3 (RMT3-23) APC | BioLegend | 119706; RRID:AB_2561656 |
| Anti-mouse CD45 (30-F11) Percp-cy5.5 | BioLegend | 103132 RRID: AB_893340 |
| Anti-mouse CD45.1 (A20) BV421 | BioLegend | 110732 BRID: AB_2562563 |
| Anti-mouse CD45.2 (104) BUV737 | BD Biosciences | 612778; RRID:AB_2870107 |
| Anti-mouse CD317 (927) Alex488 | BioLegend | 127012 RRID: AB_1953287 |
| Anti-mouse CD11c (N418) FITC | BioLegend | 117306; RRID:AB_313775 |
| Anti-mouse CD11b (M1/70) APC-Cy7 | BioLegend | 101226; RRID:AB_830642 |
| Anti-mouse CD24 (M1/69) BV711 | BioLegend | 563450; RRID:AB_2738213 |
| Anti-mouse MHC II (M5/114.15.2) BV605 | BioLegend | 107639; RRID:AB_2565894 |
| Anti-mouse F4/80 (T45-2342) BUV395 | BD Biosciences | 565614; RRID:AB_2739304 |
| Anti-mouse CD86 (GL-1) PE-Dazzle 594 | BioLegend | 105042; RRID:AB_2566409 |
| Anti-mouse CD103 (2E7) PE | BioLegend | 121406; RRID:AB_1133989 |
| Anti-mouse CD45.1 (A20) APC | BioLegend | 110714; RRID:AB_313503 |
| Anti-mouse IFN-γ (XMG1.2) PE | BioLegend | 505808; RRID:AB_315402 |

| | | |
|---|---|---|
| Anti-mouse TNF-α (MP6-XT22) APC | BioLegend | 506308; AB_315429 |
| Anti-mouse Granzyme B (QA16A02) APC | BioLegend | 372204; RRID:AB_2687028 |
| Anti-FoxP3 (150D) PE | BioLegend | 320007 AB_492981 |
| Anti-PGC-1a (D-5) PE | Santa Cruz | sc-518025 PE |
| 1E4.2.1 anti-Env antibody | Wittrup lab at MIT | N/A |
| Anti-mouse IFN-γ (XT3.11) | BioXCell | BE0055; RRID:AB_1107694 |
| Anti-mouse TNF-α (XMG1.2) | BioXCell | BE0058; RRID:AB_1107764 |
| Anti-mouse CD3ε (2C11) | BioXCell | BE0001-1 BRID:AB_1107634 |
| Anti-mouse CD28 (37.51) | BioXCell | BE0015-1 BRID:AB_1107624 |
| **Bacterial and virus strains** | | |
| 5-alpha Competent *E. coli* | New England Biolabs | C2987U |
| **Chemicals, peptides and recombinant proteins** | | |
| 1,2-distearoyl-sn-glycero-3-phosphoethanolamine-N-[maleimide (polyethylene glycol)-2000] | Layson Bio | 100220 |
| DSPE-PEG-FITC | Avanti | 810120 |
| Cyclic-di-GMP | invivogen | tlrl-nacdg |
| Resiquimod | invivogen | tlrl-r848 |
| GolgiPlug™ Protein Transport Inhibitor (containing Brefeldin A) | BD Biosciences | BDB555029 |
| Cell Stimulation Cocktail | eBioscience | 00-4970-93 |
| Protease Inhibitor Cocktail | Roche | 5892970001 |
| Recombinant murine IL-2 | Biolegend | 575408 |
| Recombinant murine IFN-γ | Peprotech | 315-05 |
| DNase I | Sigma Aldrich | 10104159001 |
| Collagenase IV | Worthington | LS004188 |
| CalPhos™ Mammalian Transfection Kit | Takara | 631312 |
| Sytox Red | Thermo Fisher | S34859 |
| Retronectin | Takara | T100B |
| TRIzol™ Reagent | Thermo Fisher | 15596018 |
| **Critical commercial assays** | | |
| NucleoSpin® Plasmid | Takara | 740588.250 |
| TrypLE™ Express Enzyme | Thermo Fisher | 12605036 |
| Gibco ACK Lysing Buffer | Thermo Fisher | A10492-01 |

| | | |
|---|---|---|
| CellTrace Violet | Thermo Fisher | C34557 |
| LIVE/DEAD™ Fixable Aqua Dead Cell Stain Kit, for 405 nm excitation | Thermo Fisher | L34966 |
| FITC Annexin V Apoptosis Detection Kit | BD Biosciences | 556547 |
| Fixation/Permeabilization Solution Kit | BD Biosciences | 554714 |
| Foxp3 / Transcription Factor Staining Buffer Set | eBioscience | 00-5523-00 |
| Mouse CD45 microbeads | Miltenyi Biotec | 130-052-301 |
| EasySep™ Mouse CD8+ T Cell Isolation Kit | Stemcell Technologies | 19853 |
| Mouse IFN-γ ELISA kit | R&D systems | DY485 |
| Mouse IL-2 ELISA kit | Invitrogen | 88-7024 |
| Mouse IFN-γ ELISPOT Kit | BD Biosciences | 551083 |
| RNeasy Micro Kit | Qiagen | 74004 |
| T-PERTM | Thermo Fisher | 78510 |
| Proteinase and phosphatase inhibitors | Thermo Fisher | 78442 |
| **Deposited data** | | |
| Bulk RNA-seq data | GEO | GSE211938 |
| sc RNA-seq data | GEO | GSE212453 |
| **Experimental Models: Cell Lines** | | |
| B16F10 cells | ATCC | CRL-6475; RRID:CVCL_0159 |
| CT-2A cells | T. Seyfried Lab at Boston college | N/A |
| MHCII$^{+}$ CT-2A cells | Generated in the Irvine lab | N/A |
| mEGFRvIII-CT-2A cells | Generated in the Irvine lab | N/A |
| ZsGreen$^{+}$ mEGFRvIII-CT-2A cells | Generated in the Irvine lab | N/A |
| mEGFRvIII-CT-2A-OVA cells | Generated in the Irvine lab | N/A |
| 293 phoenix cells | ATCC | CRL-3214 |
| B16F10-OVA cells | G. Dranoff Lab at DFCI | N/A |
| 2C TCR-58$^{-/-}$ T cell hybridoma cells | Birnbaum lab at MIT | N/A |
| 7PPG2 TCR-58$^{-/-}$ T cell hybridoma cells | Birnbaum lab at MIT | N/A |
| MC38 cells | Wittrup lab at MIT | N/A |
| TC-1 cells | ATCC | CRL-2493 |

| | | |
|---|---|---|
| TRP1$^{-/-}$ B16F10 cells | Generated in the Irvine lab | N/A |
| **Experimental Models: Organism/Strains** | | |
| C57BL/6J mice, CD45.2$^{+}$ | Jackson Laboratory | 000624; RRID:IMSR_JAX:000624 |
| C57BL/6J mice, CD45.1$^{+}$ | Jackson Laboratory | 002014; RRID:IMSR_JAX:002014 |
| Rag1$^{-/-}$ (B6.129S7-*Rag1*$^{tm1Mom}$/J) | Jackson Laboratory | 013755; RRID:IMSR_JAX:013755 |
| IFNGR1$^{-/-}$ (B6.129S7-*Ifngr1*$^{tm1Agt}$/J) | Jackson Laboratory | 003288; RRID:IMSR_JAX:00328 |
| *Batf3*$^{-/-}$ (B6.129S(C)-*Batf3*$^{tm1Kmm}$/J) | Jackson Laboratory | 013755; RRID:IMSR_JAX:013755 |

#### STAR Methods

##### Resource availability

**Materials availability.** New plasmids from this paper are available from the lead contact upon request.

**Data and code availability.** Bulk-RNA seq and single cell RNA-seq data have been deposited at GEO (GSE211938, GSE212453) and are publicly available as of the date of publication. This paper does not report original code. Any additional information required to reanalyze the data reported in this paper is available from the lead contact upon request.

##### Experimental model and subject details

**Cell line and Constructs**. B16F10 and 293 phoenix cells were obtained from ATCC. B16F10-OVA cells were a gift from Dr. Glen Dranoff at the Dana Farber Cancer Institute. TRP1$^{-/-}$ B16F10 cells were generated previously using CRIPSR[1]. The mouse CT-2A glioma cell line was kindly provided by Dr. Thomas Seyfried from Boston College. mEGFRvIII-expressing CT-2A cells were generated by lentiviral transduction of CT-2A cells with a murine version of EGFRvIII and stably selected with puromycin. ZsGreen$^{+}$ mEGFRvIII-CT-2A cells were generated by transducing mEGFRvIII-CT-2A cells with ZsGreen-expressing lentivirus and subsequent flow cytometry enrichment. mEGFRvIII-CT-2A-OVA cells were generated by transducing mEGFRvIII-CT-2A cells with PLKO-based lentivirus expressing Thy1.1-IRES-OVA (aa251-388). MHCII$^{+}$ CT-2A cells were generated by transducing CT-2A cells with lentivirus expressing CIITA (Class II Major Histocompatibility Complex Transactivator).

**Animals.** Wildtype female C57BL/6J mice (B6, CD45.2$^{+}$), CD45.1$^{+}$ congenic mice (B6.SJL-*Ptprc*$^{a}$ *Pepc*$^{b}$/BoyJ), Rag1$^{-/-}$ (B6.129S7-*Rag1*$^{tm1Mom}$/J, B6 background), IFN-γ$^{-/-}$ (B6.129S7-*Ifng*$^{tm1Ts}$/J, congenic with B6, backcrossed for at least 8 generations ), IFNGR1$^{-/-}$ (B6.129S7-*Ifngr1*$^{tm1Agt}$/J, B6 background), *Batf3*$^{-/-}$ (B6.129S(C)-*Batf3*$^{tm1Kmm}$/J, B6 background), PGC-1α-flox (B6N.129(FVB)-*Ppargc1a*$^{tm2.1Brsp}$/J), LCK-cre (B6.Cg-Tg(Lck-cre)548Jxm/J, Hemizygous), IL12rb2$^{-/-}$ (B6;129S1-

*Il12rb2$^{tm1Jm}$*/J, B6 background), IL12p40$^{-/-}$ (B6.129S1-*Il12b$^{tm1Jm}$*/J, congenic with B6, backcrossed for at least 9 generations), CD11c-cre (C57BL/6J-Tg(Itgax-cre,-EGFP)4097Ach/J, Hemizygous) and IFNGR1-flox (C57BL/6N-*Ifngr1$^{tm1.1Rds}$*/J) mice were purchased from the Jackson Laboratory. To avoid neonatal lethality caused by whole body KO of PGC-1α, T cell-specific PGC-1α KO mice were created by crossing LCK-cre mice with PGC-1α-flox mice; cre$^{+}$ F1 offspring have T cell-specific PGC-1α KO while the cre$^{-}$ F1 offspring have a wildtype phenotype and were used as donor control T cells for Fig 3F. CD11c$^{\Delta IFNGR1}$ mice were generated by crossing CD11c-cre mice with IFNGR1-flox mice, cre$^{+}$ F1 offspring are IFNGR1-deficient in CD11c$^{+}$ cells while the cre$^{-}$ F1 offspring have a wildtype phenotype and were used as control recipients in Fig. 6H. For all studies, 6-8 weeks old mice were used. All animal studies were carried out following an IACUC-approved protocol following local, state, and federal guidelines.

**Methods details**

**Cloning and constructs.** The murine EGFRvIII CAR (28z) and FITC/TA99 bispecific CAR (28z) were cloned into an MSCV retroviral vector as previously described[2]. The NFAT-IFN-γ cassette was constructed in a self-inactivating(SIN)-retroviral vector with 6xNFAT binding sites[3] upstream of the minimal IL2 promoter driving murine IFN-γ expression.

**Primary mouse T cell isolation and CAR T cell production**. For T cell activation, 6-well plates were pre-coated with 5 ml of anti-CD3 (0.5 μg/ml, Clone: 2C11) and anti-CD28 (5 μg/ml, Clone: 37.51) per well at 4°C for 18 hr. CD8$^{+}$ T cells were isolated using a negative selection kit (Stem Cell Technology), and seeded onto pre-coated 6-well plates at 5 x10$^{6}$ cells/well in 5 ml of complete medium (RPMI + penicillin/streptomycin + 10% FBS + 1x NEAA + 1x Sodium pyruvate + 1x 2-mercaptoethanol + 1x ITS [Insulin-Transferrin-Selenium, ThermoFisher]). Cells were cultured at 37°C for 48 hr without disturbance. Twenty-four hr before transduction, non-TC treated plates were coated with 15 μg/ml of retronectin (Clonetech). On day 2, cells were collected, counted and resuspended at 2x10$^{6}$ cells/ml in complete medium supplemented with 20 μg/ml of polybrene and 40 IU/mL of mIL-2. Retronectin-coated plates were blocked with 0.05% FBS containing PBS for 30 min before use. 1 ml of virus supernatant was first added into each well of the blocked retronectin plate, then 1 mL of the above cell suspension was added and mixed well by gentle shaking to reach the working concentration of polybrene at 10 μg/ml and mIL-2 at 20 IU/ml. Spin infection was carried out at 2000x*g* for 120 min at 32°C. Plates were then carefully transferred to an incubator and maintained overnight. On day 3, plates were briefly centrifuged at 1,000x*g* for 1 min, and virus-containing supernatants were carefully removed. 3 mL of fresh complete medium containing 20IU/ml of mIL-2 were then added into each well. Cells were passaged 1:2 every 12 hr with fresh complete medium containing 20IU/mL of mIL-2. Transduction efficiency was evaluated by surface staining of a c-Myc tag included in the CAR construct[2] using an anti-Myc antibody (Cell signaling, Clone:9B11) ~30 hr after transduction. If needed, CAR T cells on day 3, after flow cytometry analysis of virus transduction, could be frozen down and stored for assays at a later time. For in vivo experiments, CAR T cells were used on day 4. For *in vitro* experiments, CAR T cells were cultured till day 5.

**Virus production and transduction evaluation**. For optimal retrovirus production, 293 phoenix cells were cultured till 80% confluence, then split at 1:2 for further expansion. 24 hr later, 5.6x10$^{6}$ cells were seeded in a 10 cm dish and cultured for 16 hr till the confluency reached 70%. 30 min – 1 hr before transfection, each 10 cm dish was replenished with 10 ml pre-warmed medium. Transfection was carried out using the calcium phosphate method following the manufacturer's protocol (Clonetech). Briefly, for each transfection, 18 μg of plasmid (16.2 μg of CAR plasmid plus 1.8 μg of Eco packaging plasmid) was added to 610 μl of $ddH_2O$, followed by addition of 87 μl of 2 M $CaCl_2$. 700 μl of 2x HBS was then added in a dropwise manner with gentle vortexing. After a

10 min incubation at 25°C, the transfection mixture was gently added to phoenix cells. After 30 min incubation at 37°C, the plate was checked for the formation of fine particles, as a sign of successful transfection. The next day, old medium was removed and replenished with 8 ml of pre-warmed medium without disturbing the cells. Virus-containing supernatant was collected 36 hr later and passed through a 0.45 um filter to remove cell debris, designated as the "24hr" batch. Dishes were refilled with 10ml of fresh medium and cultured for another 24 hr to collect viruses again, designated as the "48hr" batch, this process can be repeated for another two days to collect a "72hr" batch and "96hr" batch. All virus supernatant was aliquoted and stored at -80°C. Virus transduction rate was evaluated in a 12-well format by mixing 0.5 million activated T cells with 0.5ml of viruses from each batch. Plate coating, spin infection and FACS analysis of CAR expression were carried out as described above. In the majority of experiments, the "48hr" and "72hr" batches yielded viruses that transduced T cells at 90-95% efficiency, the "24hr" and "96hr" batch viruses led to >80% transduction. Only viruses with >80% transduction rate were used for animal studies.

**Amphiphile-ligand production and vaccination**. DSPE-PEG-FITC was purchased from Avanti. Amph-pepvIII was produced as previously described[4]. Briefly, pepvIII peptides (LEEKKGNYVVTDHC) were dissolved in dimethylformamide at 10 mg/mL and mixed with 2.5 equivalents of 1,2-distearoyl-*sn*-glycero-3-phosphoethanolamine-N-[maleimide(polyethylene glycol)-2000] (Laysan Bio, Inc), 1 equivalent of tris(2-carboxyethyl)phosphine hydrochloride (Sigma), and a catalytic amount (~10ul) of triethylamine. The mixture was agitated at 25°C for 24 hr. Unconjugated peptides were removed using HPLC. Amph-pepvIII concentration was determined using nanodrop. The resulting products were lyophilized, re-dissolved in PBS and stored at -20°C. For vaccination, unless otherwise stated, mice received weekly s.c injection of 10 μg peptide equivalent of amph-pepvIII mixed with 25 μg of Cyclic-di-GMP (CDG, Invivogen) in 100 μl 1x PBS, administered 50 μl to each side at the tail base. To compare the effect of adjuvants on vaccination, 1.24 nmol lipo-CpG[4] or 10μg R848 (TLR7/8 agonist, Resiquimod [Invivogen]) was used per mouse.

**ELISPOT.** To evaluate epitope spreading, the spleen was harvested from individual mice for total T cell isolation using a $CD3^+$ T cell isolation kit (Stem Cell Technology). The day before T cell isolation, $2x10^6$ tumor cells (CT-2A, $MHCII^+$CT-2A or B16F10 cells) were seeded in a T75 flask in the presence of 100 IU of murine IFN-γ [PeproTech], and subjected to 120Gy of irradiation the next morning. Tumor cells were then trypsinized into single cell suspension using TrypLE Express (Gibco) to avoid removal of surface proteins, and washed twice with 1x PBS to remove residual IFN-γ. $4x10^5$ $CD3^+$ T cells were mixed with 25,000 irradiated tumor cells in 200 μL complete medium and seeded in a 96-well ELISPOT plate (BD) that was pre-coated with IFN-γ capture antibody (BD IFN-γ ELISPOT kit). Plates were wrapped in foil and cultured for 24hr in 37°C incubator, then developed according to the manufacturer's protocol. Plates were scanned using a CTL-ImmunoSpot Plate Reader, and data were analyzed using CTL ImmunoSpot Software.

**CAR T functionality assay**. The functionality of WT, IFN-$\gamma^{-/-}$, IFNGR1$^{-/-}$ or NFAT-IFNγ CAR T cells was assessed by co-coculturing with EGFRvIII-CT2A cells in 96-well flat-bottom plates. Unless otherwise stated, $1x10^5$ CAR T cells were mixed with $1x10^4$ target cells in a total volume of 200 μl complete medium containing 20IU/ml of mIL-2. After 6 hr co-culture, cells were resuspended by vigorous pipetting, transferred to a U-bottom plate, and pelleted at 2,000x*g* for 5 min. The supernatant was saved for ELISA following the manufacturer's protocol (Mouse IFN-γ Duo set, R&D systems). Cells were stained with anti-CD45 and anti-CD8α for 20 min on ice, and resuspended in flow cytometry buffer with 1x SYTOX Red (Thermo Fisher) for flow analysis. Dead tumor cells were gated as CD8⁻CD45⁻ SYTOX RED⁺ population. IFN-γ ELISAs were performed following the manufacturer's protocol.

**P15E antigen and Env protein detection**. Env protein expression on CT-2A cell surface was monitored using flow cytometry and staining with 1E4.2.1 anti-Env antibody as previously described[5] (Wittrup lab). The presentation of Env antigen p15E on CT-2A cells were assessed by co-culturing IFN-γ-treated CT-2A cells with a $58^{-/-}$ T cell hybridoma cell line expressing a p15E-specific TCR 7PPG-2 and monitoring T cell activation using mouse IL-2 ELISA (Invitrogen) as previously described[2,6]. A $58^{-/-}$hybridoma cells expressing an irrelevant 2C TCR were included as negative control. TC-1 cells and MC38 cells were included as negative control and positive control of ENV/p15E expression, respectively.

**Secondary transplantation study.** To evaluate the qualitative anti-tumor activity of CAR T cells, 10 million $CD45.1^+$ donor CAR T cells were i.v. infused to lymphodepleted $CD45.2^+$ recipients (500cGy sublethal irradiation) followed 24hr later by a single dose of amph-pepVIII vaccination or mock vaccination with PBS. Seven days later, mice were euthanized and spleens were harvested and combined for each group for total T cell isolation using a modified pan-T cell negative selection protocol. Briefly, total splenocytes were stained with a pan-T cell isolation cocktail (Stem Cell Technology) plus 1:500 dilution of biotinylated anti-CD45.2 antibody (0.5mg/ml, Stem Cell Technology). Negative selection was performed following the same downstream procedures as listed in the manufacture's protocol to obtain untouched vaccine-boosted CD45.1 CAR T cells. Immediately after isolation, $8x10^6$ of CAR T cells from either mock or vaccine-treated groups were adoptively transferred to secondary recipients bearing ~25 $mm^2$ EGFRvIII-CT-2A tumors that had been lymphodepleted the day before, followed by periodic monitoring of tumor growth and animal survival.

**Luminex assay.** EGFRvIII-CT-2A tumor-bearing C57BL/6 mice received lymphodepletion followed by adoptive transfer of either WT or IFN-$\gamma^{-/-}$ CAR T cells plus a single dose of vaccination. Mice were euthanized and tumors isolated at day 7 post vaccination. Tumors were weighted, cut using a razor blade into small pieces and dounced to generate tumor homogenate in tissue protein extraction buffer (T-PERTM, Thermo Fisher Scientific, cat. no. 78510) in the presence of 1% proteinase and phosphatase inhibitors (Thermo Fisher Scientific, cat. no. 78442). The lysates were incubated at 4°C for 30 min with slow rotation followed by top-speed centrifugation to remove debris. The supernatants were transferred to a clean tube and stored at -80°C. Part of the samples were subjected to Luminex analysis using a Mouse Cytokine 32-Plex panel analysis at Eve Technology.

**Bulk RNA-sequencing for CAR T characterization**. EGFRvIII-CT-2A tumor-bearing $CD45.2^+$ mice were treated with $CD45.1^+$ CAR T cells and mock (PBS) or amph-pepVIII vaccination. 7 days later, mice were euthanized to harvest spleens and tumors. Total splenic T cells were isolated using the pan-T cell isolation kit and stained with anti-CD8α, anti-CD4, anti-CD45.1 and 7AAD for flow sorting. $5x10^4$ $CD45.1^+$ CAR T cells were directly sorted into Trizol. For intratumoral CAR T isolation, tumors were cut into 1-2 $mm^2$ pieces using razor blades, placed in 1.5ml or 5ml tubes (depending on tumor size) and digested (2 mg/ml Collagenase IV [Worthington], 0.1mg/ml of DNAse I [Sigma], and 1% of TrypLE [ThermoFisher] in 1xRPMI) for 30 min on a rotator at 37°C. Intratumoral T cells were enriched using mouse CD4/CD8 (TIL) MicroBeads (Miltenyi), stained, and sorted into Trizol as above. The total number of sorted $CD45.1^+$ CAR T cells from tumors ranged from $6x10^3$ to $5x10^4$ per sample. Total RNA was isolated using the Reasy Micro kit (Qiagen). Samples were submitted to the BioMicro center at MIT for library construction and sequencing. Bulk RNA-sequencing data was analyzed with the help from the bioinformatics core at the Koch Institute. Briefly, paired-end RNA-seq data was used to quantify transcripts from the mm10 mouse assembly with the Ensembl version 100 annotation using Salmon version 1.2.1[7]. Gene level summaries for were prepared using tximport version 1.16.0[8]. running under R version 4.0.0 (https://www.R-project.org). Differential expression analysis was performed using DESeq2 version 1.28.1[9,10] and differentially expressed genes were defined as those having an absolute apeglm[11] log2 fold change greater than 1 and an adjusted p-value less than 0.05. Data parsing and some visualizations were carried out using Tibco Spotfire Analyst 7.6.1. Mouse genes were mapped to human orthologs using Mouse Genome Informatics (http://www.informatics.jax.org/) orthology report. Preranked GSEA[12] was run using javaGSEA version 4.0.3 for gene sets from MSigDB version 7.1[13]. Preranked GSEA for custom mouse gene sets was run with javaGSEA version 4.1.0

**Seq-Well Single cell RNA-sequencing to profile AS in intratumoral T cells**. Tumors were digested and tumor-infiltrating lymphocytes enriched as described above for bulk RNAseq. Enriched TILs from individual mice were first labeled with Total-seq A anti-mouse hashing antibodies (Biolegend) and washed 2x in flow cytometry buffer. Samples from the same group were then combined and stained with the same surface staining antibody cocktail. Endogenous $CD45.2^+$ CD4 and CD8 T cells were sorted collectively into 1x RPMI +10%FBS, 2-5x$10^4$ total T cells were obtained for each group. Cells were pelleted at 1000xg for 5min, resuspended in 1xRPMI at 20,000 cells per 200µl and then processed for scRNA-seq using the Seq-Well platform with second strand chemistry, as previously described[14]. Whole transcriptome libraries were barcoded and amplified using the Nextera XT kit (Illumina) and were sequenced on a Novaseq 6000 (Illumina). Hashtag oligo libraries were amplified as described previously[15] and were sequenced on a Nextseq 550.

**Processing of single cell hashing data.** Cell hashing data was aligned to HTO barcodes using CITE-seq-Count v1.4.2 (https://zenodo.org/badge/latestdoi/99617772). To establish thresholds for positivity for each HTO barcode, we first performed centered log-ratio normalization of the HTO matrix and then performed k-medoids clustering with k=5 (one for each HTO). This produced consistently five clusters, each dominated by one of the 5 barcodes. For each cluster, we first identified the HTO barcode that was dominant in that cluster. We then considered the threshold to be the lowest value for that HTO barcode among the cells classified in that cluster. To account for the scenario in which this value was substantially lower than the rest of the values in the cluster, we used Grubbs' test to determine whether this threshold was statistically an outlier relative to the rest of the cluster. If the lower bound was determined to be an outlier at p=0.05, it was removed from the cluster, and the next lowest value was used as the new threshold. This procedure was iteratively applied until the lowest value in the cluster was no longer considered an outlier at p=0.05. Cells were then determined to be "positive" or "negative" for each HTO barcode based on these thresholds. HTO thresholds were examined and manually adjusted if necessary. Cells that were positive for multiple HTOs (doublets) or were negative for all HTOs were excluded from downstream analysis. To account for differences in sequencing depth between samples, these steps were performed separately for each Seq-Well array that was processed.

**scRNA-seq data processing and visualization**. Raw read processing of scRNA-seq reads was performed as previously described[16]. Briefly, reads were aligned to the mm10 reference genome and collapsed by cell barcode and unique molecular identifier (UMI). Then, cells with less than 500 unique genes detected and genes detected in fewer than 5 cells were filtered out, and the data for each cell was log-normalized to account for library size. Genes with log-mean expression values greater than 0.1 and a dispersion of greater than 1 were selected as variable genes, and the ScaleData function in Seurat was used to regress out the number of UMI and percentage of mitochondrial genes in each cell. Principal components analysis was performed. The number of principal components used for visualization was determined by examination of the elbow plot, and two-dimensional embeddings were generated using uniform manifold approximation and projection (UMAP). Clusters were determined using Louvain clustering, as implemented in the FindClusters function in Seurat, and clusters that contained activated T cells were selected for further analysis. These cells were reprocessed with the same processing and clustering steps described above. DEG analysis was performed for each cluster and between indicated cell populations using the FindMarkers function.

**Paired single-cell TCR sequencing and analysis**. Paired TCR sequencing and read alignment was performed as previously described[17]. Briefly, whole transcriptome amplification product from each single-cell library was enriched for TCR transcripts using biotinylated *Tcrb* and *Tcra* probes and magnetic streptavidin beads. The enrichment product was further amplified using V-region primers and Nextera sequencing handles, and the resulting libraries were sequenced on an Illumina Novaseq 6000. Processing of reads was performed using the Immcantation software suite[18,19]. Briefly, reads were aggregated by cell barcode and UMI, and UMI with under 5 reads were discarded. ClusterSets.py was used to divide sequences for each UMI into sets of similar sequences. Only sets of sequences that comprised greater than 90% of the sequences obtained for that UMI were considered further. Consensus sequences for each UMI were determined using the BuildConsensus.py function. Consensus sequences were then mapped against TCRV and TCRJ IMGT references sequences with IgBlast. Sequences for which a CDR3 sequence could not be unambiguously determined were discarded. UMI for consensus sequences were corrected using a directional UMI collapse, as implemented in UMI-tools[20]. TCR sequences were then mapped to single cell transcriptomes by matching cell barcodes. If multiple *Tcra* or *Tcrb* sequences were detected for a single cell barcode, then the corresponding sequence with the highest number of UMI and raw reads was retained. TCR data for p15E tetramer-sorted CD8+ T cells was obtained from Grace et al(Grace 2022). Using this data, we defined high-confidence p15E-specific Tcrb and Tcra CDR3 amino acid sequences as sequences that were detected in more than one cell and for which greater than 80% of total sequences recovered were in the tetramer-positive fraction. Using this set of sequences as a reference, we defined likely p15E-specific clonotypes in our sequencing of TIL from CAR T and CAR T-Vax treated mice as clonotypes that utilized either one of these Tcrb or Tcra amino acid sequences or utilized the Tcra motif "DYSNNRLT", which was strongly implicated in the recognition of the p15E epitope by Grace et al. To define a cytotoxicity score for each CD8+ T cell in our single-cell sequencing data, we utilized the AddModuleScore function in Seurat using the following genes as a signature: Gzma, Gzmb, Gzmc, Gzmd, Gzme, Gzmf, Gzmg, Gzmk, Gzmm. Single T cells for which neither a Tcrb or Tcra sequence were recovered were excluded from this analysis.

**Phenotyping of immune cells in peripheral blood, lymph nodes, and tumors**. Peripheral blood (PB) was collected via retro-orbital bleeding. 50-100 μl PB (lymphodepleted mice) was processed in ACK lysis buffer twice (3-5min the 1st time till all RBCs were lysed followed by centrifugation at 1000xg for 5min, decant, resuspend in 200ul ACK and spin again), immediately after spin the 2nd time, instead of decanting, RBC debris from each well was carefully removed by vacuuming in a circular motion without touching the center of the pellet. Lymph nodes (LNs) were placed in a 5ml flow cytometry tube with a 70 μM cell strainer cap and smashed through with the rubber end of a 1ml syringe plunger with frequent addition of flow cytometry buffer. Dissociated LN cells were pelleted and transferred to a 96 well U-bottom plate for further analysis. EGFRvIII-CT-2A tumors from mice receiving CAR T or CAR T-vax therapy were surgically removed and weighed and dissociated into single cell suspension using enzyme digestion as described above. Single cell suspensions were obtained by passing tumors through a 70 μM cell strainer with a 1 ml syringe plunger. Cells were pelleted and resuspended with 100 μl of FACS buffer per 100 mg tumor. For immunophenotyping analysis, PBMCs or lymph node cell suspensions were pelleted in a 96 well U-bottom plate and stained with desired antibody cocktails at a 1:200 dilution for $CD4^+$ T cells, $CD8^+$ T cells, Tregs, B cells, $CD103^+$ cDC1, $CD11b^+$ cDC2, pDCs.

For intracellular cytokine staining (ICS) analysis, 50-75 μl of the above tumor cell suspension was pelleted in 96 well U-bottom plates and directly resuspended in RPMI1640 with 10% FBS plus 1x Golgi plug and 1x cell stimulation cocktail (Thermo) for 6 hr at 37°C. Cells were then pelleted at 1000xg for 5 min and washed once with PBS, then stained with live/dead aqua for 15 min in the dark at 25°C. Cells were pelleted again, surface stained with desired antibody cocktail for ~20 min on ice followed by 1x wash with flow cytometry buffer. Cells were resuspended in 75 μl of BD Fix/Perm and kept at 4°C for 15 min, then washed once by directly adding 200 μl 1x Perm/Wash. The pellet was resuspended in 50 μl of cytokine antibody cocktail (IFN-γ at 1:100, TNF-α at 1:100, and Granzyme B at 1:100) pre-diluted in 1x Perm/Wash buffer, 30 min on ice, then washed once with 1x Perm/Wash buffer and resuspended in 1x flow cytometry buffer for analysis immediately or kept at 4°C for FACS analysis the next day. For FoxP3 or PGC-1α staining, 50-75μl of the above cell suspension was pelleted and processed using a FoxP3 staining kit (Thermo Fisher) according to the manufacturer's instructions.

For tetramer staining, PBMCs or tumor suspensions were stained with 50 μl of SIINFEKL-Tetramer (PE conjugate) plus Fc block at a 1:50 dilution for 30 min at room temperature in the dark, followed by mixing with a pre-made 50 μl cocktail of the remaining surface antibodies (1:50 dilution), 20min on ice kept from light. Then cells were washed twice for flow analysis.

**CAR T-vax therapy in solid tumor models**. In the EGFRvIII-CT-2A mouse glioblastoma model, unless otherwise stated, $5x10^6$ EGFRvIII-CT-2A cells were injected into the right flank of recipient mice in 50 μl saline and allowed to establish palpable tumors ~25 $mm^2$ in size at day 6. Lymphodepletion was carried out using 500 cGy sublethal irradiation, mice were then randomly allocated into each group. $10x10^6$ CAR T cells from mice with the desired background were i.v. infused via the tail vein into recipient mice followed by weekly s.c. immunization with amph-pepvIII vaccine (10 μg amph-pepvIII, 25 μg CDG in 100μl PBS) or PBS alone. For consistency, $Rag1^{-/-}$ mice were also subjected to the same lymphodepletion preconditioning. For experiments involving cytokine blockade, unless otherwise stated, anti-IFN-γ (BioXcell) was administered i.p. at 200 μg per mouse every three days, anti-TNF-α (BioXcell) was administered i.p. at 300 μg per mouse every two days.

For the mixed tumor studies, each mouse was inoculated in the right flank with $5x10^6$ EGFRvIII-CT-2A cells and WT CT-2A cells mixed at pre-defined ratios (100:0, 90:10, 80:20, 50:50, 25:75, 0:100). Six days later, when the tumors reached ~25$mm^2$, mice were subjected to lymphodepletion and adoptive transfer of $10x10^6$ EGFRvIII CAR T cells, followed 24hr later with weekly vaccination.

In the B16F10-OVA mouse melanoma model, B16F10-OVA tumors were established by s.c injection of $1x10^6$ B16F10 cells into the right flank of C57BL/6 recipient mice in 50 μl saline. Mice were lymphodepleted with 500 cGy sublethal irradiation at day 6, and received $10x10^6$ CD45.1 FITC/TRP1 bispecific CAR T cells at day 5, followed by two weekly amph-FITC immunizations (10nmol amph-FITC, 25 μg CDG in 100μl PBS).

**Statistics, selection of animals and justification of sample size**. Statistical analyses were performed using GraphPad Prism 8. All values and error bars are shown as mean ± 95% CI (confidence interval). Animal survival was analyzed using Log-rank (Mantel-Cox) test. All pair-wise comparisons were analyzed by student's t-test. Multi-group comparisons was carried out using one-way ANOVA with Tukey's multiple comparisons test. Experiments that involved repeated measures over a time course, such as tumor growth, were analyzed using a RM (repeated measures) two-way ANOVA based on a general linear model (GLM). The RM design included factors for time, treatment and their interaction. Tukey's multiple comparisons test was carried out for the main treatment effect. P-values are adjusted to account for multiple comparisons in both one-way ANOVA, and RM two-way ANOVA. For all animal experiments, 6-8 week old female C57BL/6 were used. At this age, mice have developed a mature immune system, thus ideal for evaluating immunomodulating therapies. We determined the size of samples for experiments involving either quantitative or qualitative data as previously reported[21]. Based on our previous experience with the animal models and as reported by others[1,22,23], we consider the CAR T-vax therapy as significant if it increases the survival of animals up to 100% within 4 weeks, and we need >=5 animals per group to achieve this goal with 95% confidence interval and at 80% power.

**Supplemental Table 1**
Differential gene expression for the single cell clusters defined in Figure 2C-D.

**Supplemental Table 2**
List of genes differentially expressed in $CD8^+$ and $CD4^+$ TILs at day 7 and day 14 between CAR T and CAR T-vax treatment groups in Figure 2 and Figure S2.

**Supplemental Table 3**
List of T cell clones with predicted p15E-antigen specificity in T cells from Figure 2.

**Supplemental Table 4**
List of genes differentially expressed in splenic CAR T cells treated with or without vaccination from Figure 4B.

**Supplemental Table 5**
List of genes differentially expressed in $CD8^+$ and $CD4^+$ TILs at day 14 from mice receiving CAR T-vax with or without IFNγ blockade in Figure S5D-E.

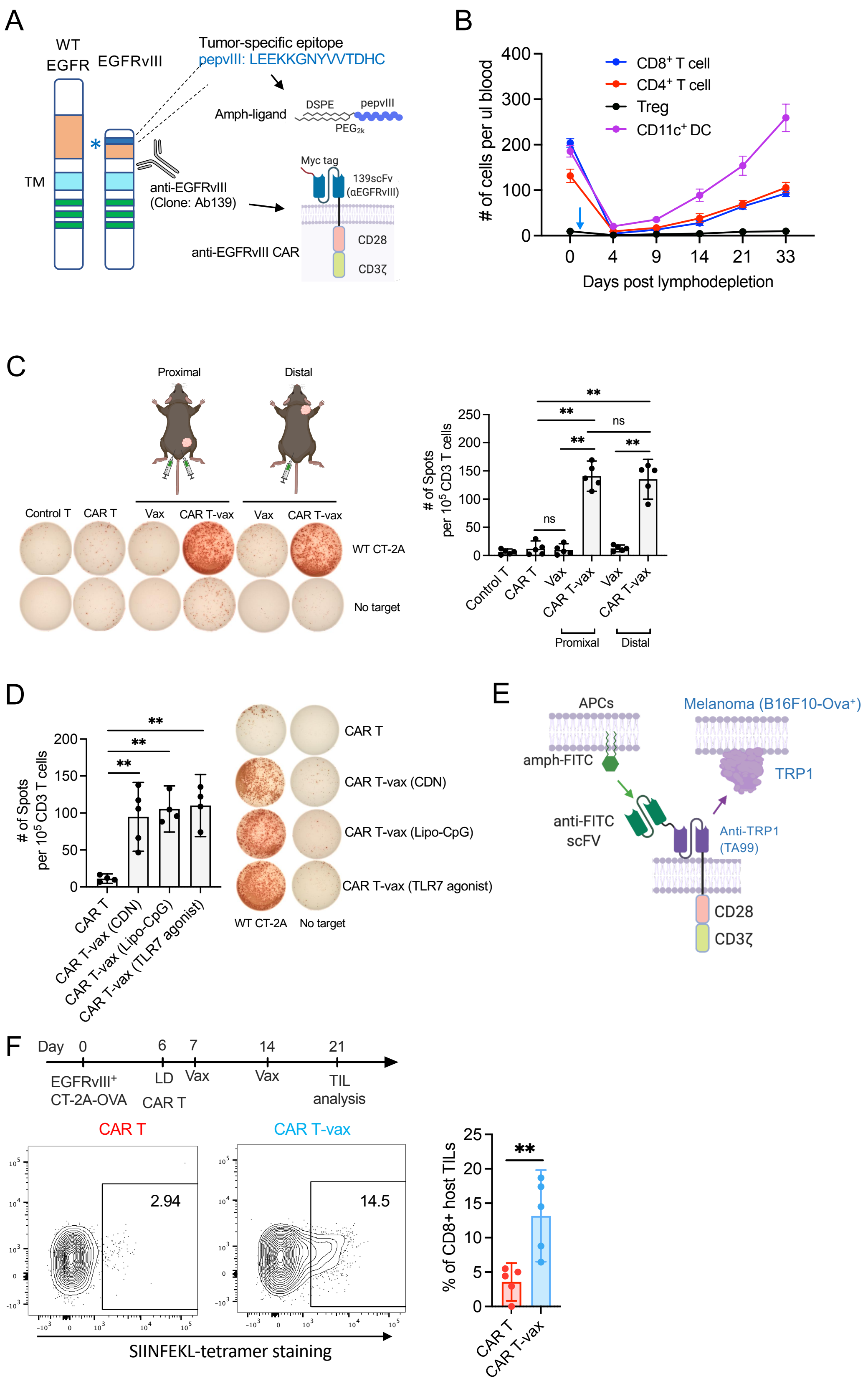
A
WT EGFR
EGFRvIII
Tumor-specific epitope
pepvIII: LEEKKGNYVVTDHC
Amph-ligand
DSPE
pepvIII
PEG2k
TM
anti-EGFRvIII (Clone: Ab139)
Myc tag
139scFv (αEGFRvIII)
CD28
CD3ζ
anti-EGFRvIII CAR
B
### of cells per ul blood
CD8+ T cell
CD4+ T cell
Treg
CD11c+ DC
Days post lymphodepletion
C
Proximal
Distal
Control T
CAR T
Vax
CAR T-vax
WT CT-2A
No target
### of Spots per 10^5 CD3 T cells
ns
**
Promixal
Distal
D
CAR T
CAR T-vax (CDN)
CAR T-vax (Lipo-CpG)
CAR T-vax (TLR7 agonist)
WT CT-2A
No target
E
APCs
Melanoma (B16F10-Ova+)
amph-FITC
TRP1
anti-FITC scFV
Anti-TRP1 (TA99)
CD28
CD3ζ
F
Day 0 6 7 14 21
EGFRvIII+ CT-2A-OVA
LD
CAR T
Vax
Vax
TIL analysis
CAR T
CAR T-vax
2.94
14.5
SIINFEKL-tetramer staining
% of CD8+ host TILs
**

**Figure S1. Delineation of factors contributing to CAR T-vax therapy induced antigen spreading.**

(A) Schematic of the development of anti-EGFRvIII CAR and amph-pepvIII vaccine.

(B) Kinetics of the recovery of T cell and DC populations post sublethal irradiation (500 cGy) in C57BL/6 mice (n = 5 animals/group). Day 0 denotes the baseline level prior to irradiation. Arrow indicates day of irradiation. Shown is one representative of at least three independent experiments.

(C) Impact of tumor location and site of vaccination on the magnitude of CAR T-vax induced antigen spreading. C57BL/6 mice (*n*=5 animals/group) bearing $EGFRvIII^{+}$CT-2A tumors received lymphodepletion (LD) followed by treatment with control (untransduced) T-cells or CAR T-vax following the same timeline as in Fig 1C. IFN-γ ELISPOT was assayed for splenic T cells isolated on day 21 and stimulated with irradiated EGFRvIII-negative CT-2A cells. Shown are representative ELISPOT well images and quantitative ELISPOT data from one representative of two independent experiments.

(D) Impact of adjuvants on eliciting CAR T-vax induced antigen spreading. $EGFRvIII^{+}$CT-2A tumor-bearing C57BL/6 mice (*n*=5 animals/group) received lymphodepletion (LD) and subsequent treatment with CAR-T in the absence or presence of amph-pepvIII vaccine formulated with different adjuvants administered following the same timeline as in Fig 1C. Shown is IFN-γ ELISPOT monitoring endogenous T-cell priming across various conditions at day 21 as in (C).

(E) Schematic of CAR T-vax therapy using a combination of amph-FITC vaccine and FITC/TA99 bispecific CAR T cells.

(F) Tumor antigen specificity of endogenous TILs in mice receiving CAR T vs CAR T-vax therapy. C57BL/6 mice (*n*=5 animals/group) bearing OVA-expressing EGFRvIII-CT-2A tumors received lymphodepletion and CAR T transfer followed by two weekly vaccinations. Upper panel, experimental timeline. Lower panel, representative flow cytometry plots showing SIINFEKL-tetramer staining of endogenous $CD8^{+}$ T cells within tumors isolated from mice treated with CAR T or CAR T-vax therapy on day 21.

In Panel C-D, data shown are mean ± 95% CI. **, p<0.01; ns, not significant by one-way ANOVA with Tukey’s post-test. In panel F, data shown are mean ± 95% CI.*, p<0.05; ns, not significant by Student's *t*-test.

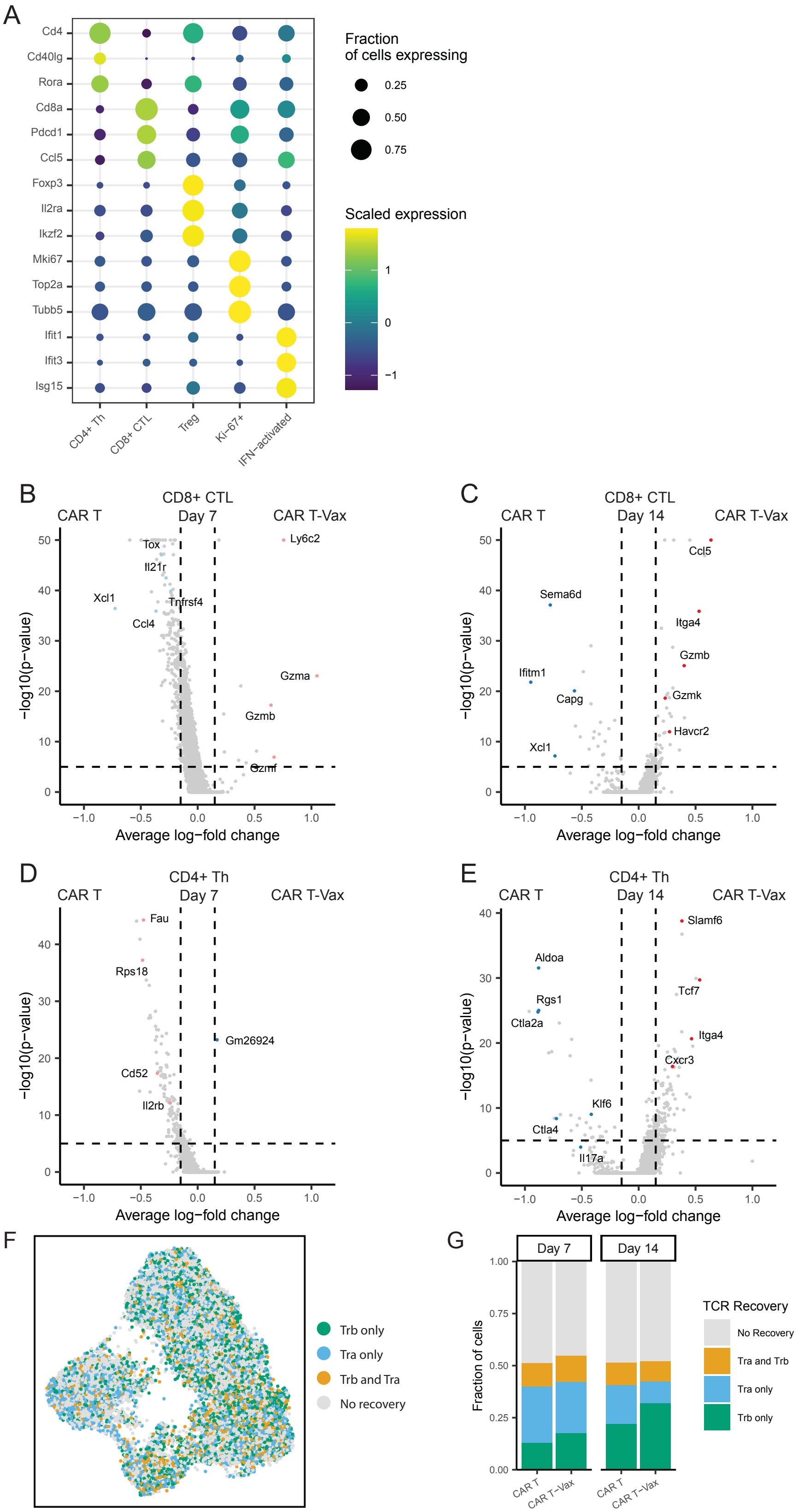

A
Cd4
Cd40lg
Rora
Cd8a
Pdcd1
Ccl5
Foxp3
Il2ra
Ikzf2
Mki67
Top2a
Tubb5
Ifit1
Ifit3
Isg15
CD4+ Th
CD8+ CTL
Treg
Ki-67+
IFN−activated
Fraction of cells expressing
0.25
0.50
0.75
Scaled expression
1
0
−1
B
CD8+ CTL Day 7
CAR T
CAR T-Vax
Tox
Il21r
Xcl1
Tnfrsf4
Ccl4
Ly6c2
Gzma
Gzmb
Gzmf
−log10(p−value)
Average log−fold change
C
CD8+ CTL Day 14
CAR T
CAR T-Vax
Ccl5
Sema6d
Itga4
Gzmb
Ifitm1
Capg
Gzmk
Havcr2
Xcl1
−log10(p−value)
Average log−fold change
D
CD4+ Th Day 7
CAR T
CAR T-Vax
Fau
Rps18
Gm26924
Cd52
Il2rb
−log10(p−value)
Average log−fold change
E
CD4+ Th Day 14
CAR T
CAR T-Vax
Slamf6
Aldoa
Tcf7
Rgs1
Ctla2a
Itga4
Cxcr3
Klf6
Ctla4
Il17a
−log10(p−value)
Average log−fold change
F
Trb only
Tra only
Trb and Tra
No recovery
G
Day 7
Day 14
Fraction of cells
CAR T
CAR T−Vax
TCR Recovery
No Recovery
Tra and Trb
Tra only
Trb only

**Figure S2. Single cell phenotyping and differential gene expression analysis**

(A) Dot plot showing differential expression of selected genes in different T cell subtypes as a result of vaccine boosting of CAR T cells in Fig 2.

(B) Volcano plot showing differential gene expression in intratumoral $CD8^+$ CTLs between CAR T-vax vs. CAR-T alone group on day 7 in Fig 2.

(C) Volcano plot showing differential gene expression in intratumoral $CD8^+$ CTLs between CAR T-vax and CAR-T alone group on day 14 in Fig 2.

(D) Volcano plot showing differential gene expression in intratumoral $CD4^+$ Th cells between CAR T-vax and CAR-T alone group on day 7 in Fig 2.

(E) Volcano plot showing differential gene expression in intratumoral $CD4^+$ Th cells between CAR T-vax and CAR-T alone group on day 14 in Fig 2.

(F) UMAP of T cells with detectable TCR alpha, beta or both chains. TCR data was extracted from the scRNA-seq data of Fig 2.

(G) Stacked charts showing proportions of T cells with detectable TCR alpha, beta or both chains on day 7 and day 14, respectively.

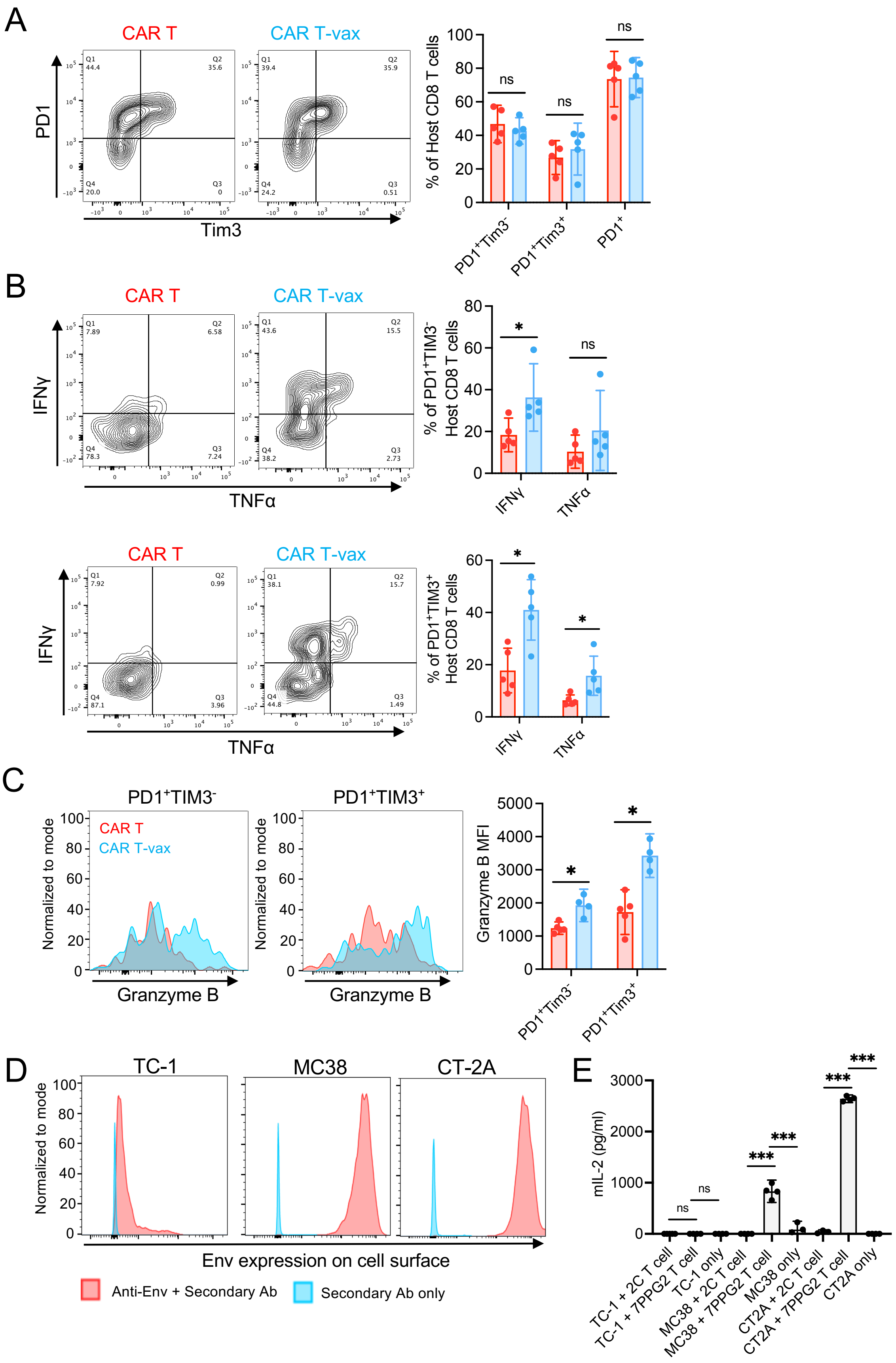

A
CAR T
CAR T-vax
PD1
Tim3
% of Host CD8 T cells
ns
PD1+Tim3-
PD1+Tim3+
PD1+
B
CAR T
CAR T-vax
IFNγ
TNFα
% of PD1+TIM3- Host CD8 T cells
% of PD1+TIM3+ Host CD8 T cells
C
PD1+TIM3-
PD1+TIM3+
Normalized to mode
Granzyme B
Granzyme B MFI
D
TC-1
MC38
CT-2A
Env expression on cell surface
Anti-Env + Secondary Ab
Secondary Ab only
E
mIL-2 (pg/ml)
TC-1 + 2C T cell
TC-1 + 7PPG2 T cell
TC-1 only
MC38 + 2C T cell
MC38 + 7PPG2 T cell
MC38 only
CT2A + 2C T cell
CT2A + 7PPG2 T cell
CT2A only

F

| Clone Name | Clone size | TRAV | CDR3α | TRAJ | TRBV | CDR3β | TRBJ |
|---|---|---|---|---|---|---|---|
| 7PEQ27 | 66 | TRAV7-2 | CAASDYSNNRLT | TRAJ7 | TRBV16 | CASSLELGGPEQYF | TRBJ2-7 |
| 7LEQ27 | 52 | TRAV3D-3 | CAVTPDYSNNRLT | TRAJ7 | TRBV16 | CASSLELGGLEQYF | TRBJ2-7 |
| 7QTG27 | 42 | TRAV8D-2 | CATDYSNNRLT | TRAJ7 | TRBV12-1 | CASSQTGGPSYEQYF | TRBJ2-7 |
| 7LEV24 | 41 | TRAV6-6 | CALLPADYSNNRLT | TRAJ7 | TRBV16 | CASSLEVGQGYSQNTLYF | TRBJ2-4 |
| 7REQ27 | 31 | TRAV7-2 | CAGSDYSNNRLT | TRAJ7 | TRBV16 | CASSLELGGREQYF | TRBJ2-7 |
| 27LEG27 | 27 | TRAV6-4 | CALANTNTGKLTF | TRAJ27 | TRBV16 | CASSLELGGLEQYF | TRBJ2-7 |
| 7LRG15 | 15 | TRAV13D-1 | CALGDYSNNRLT | TRAJ7 | TRBV3 | CASSLRGGDQAPLF | TRBJ1-5 |
| 7QEG11 | 14 | TRAV5D-4 | CAAKDYSNNRLT | TRAJ7 | TRBV2 | CASSQEGANTEVFF | TRBJ1-1 |
| 7QTG25 | 13 | TRAV9N-3 | CAVFPDYSNNRLT | TRAJ7 | TRBV2 | CASSQTGGGDTQYI | TRBJ2-5 |
| N7LEQ27 | 11 | TRAV7-2 | CAANDYSNNRLT | TRAJ7 | TRBV16 | CASSLELGGLEQYF | TRBJ2-7 |

G

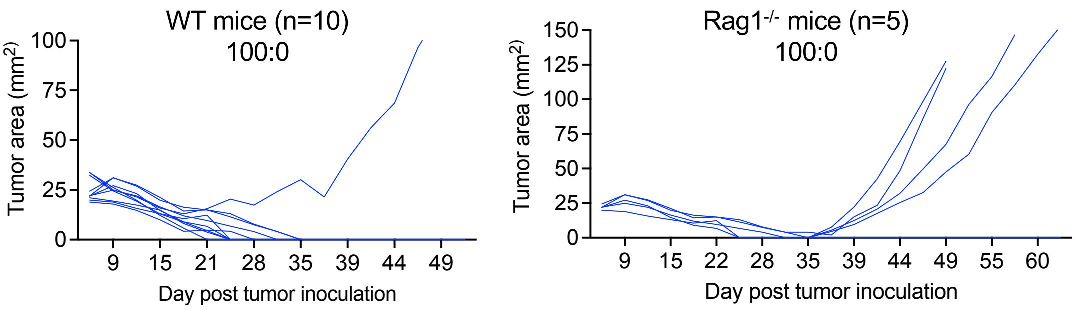


**Figure S3. Characterization of the phenotype and polyfunctionality of intratumoral endogenous T cells** and their contributions to long-term tumor control.

(A) CD45.2$^+$ C57BL/6 mice bearing EGFRvIII$^+$CT-2A tumors were treated by CD45.1$^+$ EGFRvIII-CAR T ± vax as in Fig 2B. On day 7 post therapy, tumor-infiltrating endogenous CD8$^+$ T cells were analyzed for checkpoint marker (PD1, Tim3) expression by flow cytometry. Shown is one representative of two independent experiments.

(B) Cytokine polyfunctionality in PD1$^+$TIM3$^-$ and PD1$^+$TIM3$^+$ tumor-infiltrating endogenous CD45.2$^+$ CD8$^+$ T cells from mice in Fig. S4A. Tumors were dissociated into single cell suspensions on day 7 post therapy and cultured in the presence of 1x cell stimulation cocktail and Golgi plug for 6 hours followed by intracellular cytokine staining and flow cytometry analysis.

(C) Granzyme B expression in PD1$^+$TIM3$^-$ and PD1$^+$TIM3$^+$ tumor-infiltrating endogenous CD45.2$^+$ CD8$^+$ T cells from mice in Fig. S4A and analyzed via intracellular staining as in Fig. S4B.

(D) Histogram showing retroviral Env expression on CT-2A cells, MC38 cells and TC-1 cells. Cells were stained with anti-Env primary antibody followed by anti-IgG isotype secondary antibody. MC38 and TC-1 cells were included as positive and negative controls[5], respectively.

(E) mouse IL-2 ELISA showing p15E antigen expression in CT-2A cells as detected by T cells. CT-2A cells, MC38 cells and TC-1 cells were pre-treated with IFN-γ and co-cultured with 58 T-cell hybridoma expressing an irrelevant 2C TCR or a p15E-reactive 7PPG2 TCR at 1:1 E:T ratio for 24hr. The supernatant was collected for ELISA.

(F) CDR3 sequences and gene usage for top-ranked p15E-specific T cell clones based on the CDR3 sequences and consensus motifs reported in Grace et al (Grace 2022). The consensus motifs were underlined in CDR3α or CDR3β sequences.

(G) Long-term growth of tumors in WT ($n$=10 animals/group) or Rag1$^{-/-}$ mice ($n$=5 animals/group) in Fig 3D. Only the EGFRvIII$^+$ CT-2A and EGFRvIII$^-$ CT-2A 100:0 ratio group is shown here.

In panel A-C, $n$=5 animals/group, data shown are mean ± 95% CI. *, $p<0.05$; ns, not significant by Student's $t$-test. In panel D-E, n=4 biol. replicates/group, data shown are mean ± 95% CI.***, $p<0.001$; ns, not significant by one-way ANOVA with Tukey's post-test.

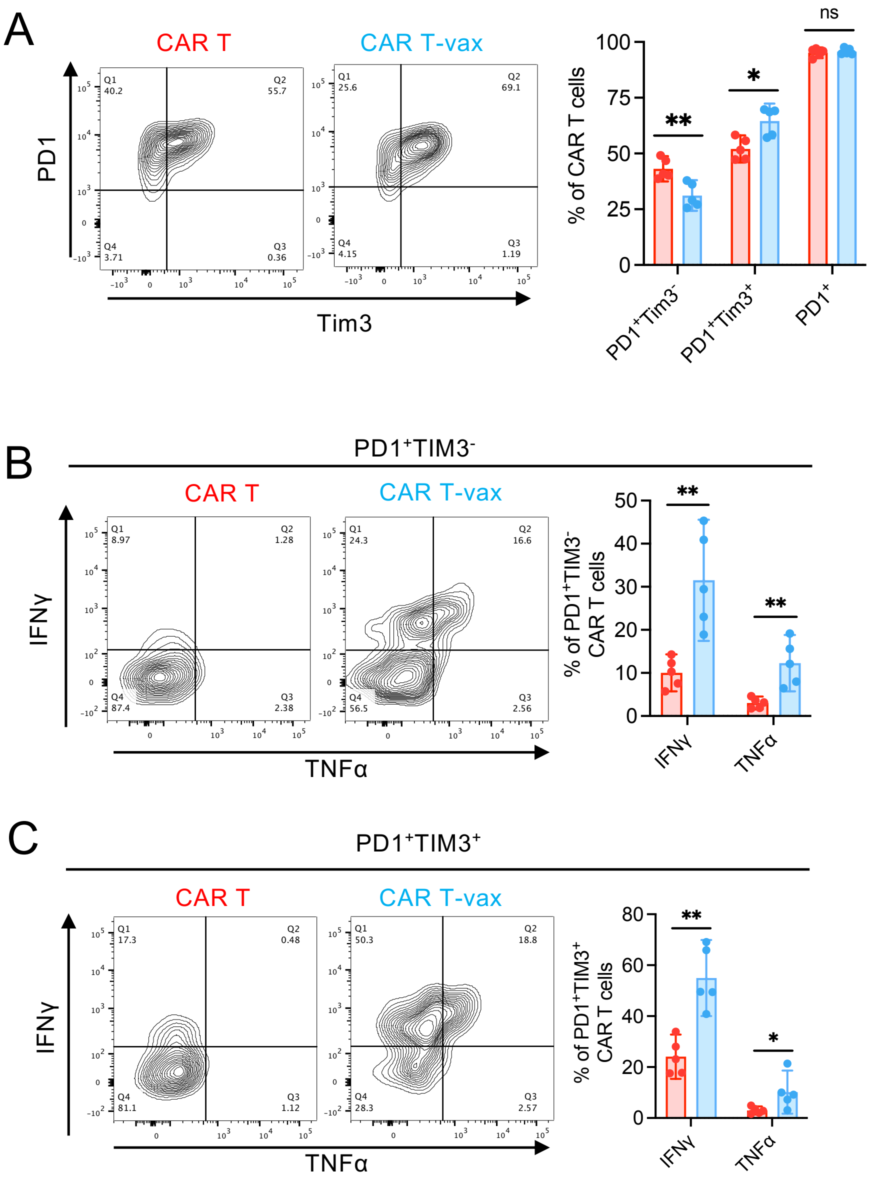


**Figure S4. Characterization of the phenotype and polyfunctionality of intratumoral CAR T cells.**

(A) CD45.2$^+$ C57BL/6 mice bearing EGFRvIII$^+$CT-2A tumors were treated by CD45.1$^+$ EGFRvIII-CAR T ± vax as in Fig 2B. On day 7 post therapy, tumor-infiltrating CAR T cells were analyzed for checkpoint marker (PD1, Tim3) expression by flow cytometry. Shown is one representative of two independent experiments.

(B-C) Cytokine polyfunctionality in PD1$^+$TIM3$^-$ (B) and PD1$^+$TIM3$^+$ (C) tumor-infiltrating CD45.1$^+$ CAR T cells from mice in Fig. S6A. Tumors were dissociated into single cell suspensions on day 7 post therapy and cultured in the presence of 1x cell stimulation cocktail and Golgi plug for 6 hours followed by intracellular cytokine staining and flow cytometry analysis.

Throughout, *n*=5 animals/group, error bars are mean ± 95% CI, **p<0.01, *p<0.05, n.s not significant by Student's t-test.

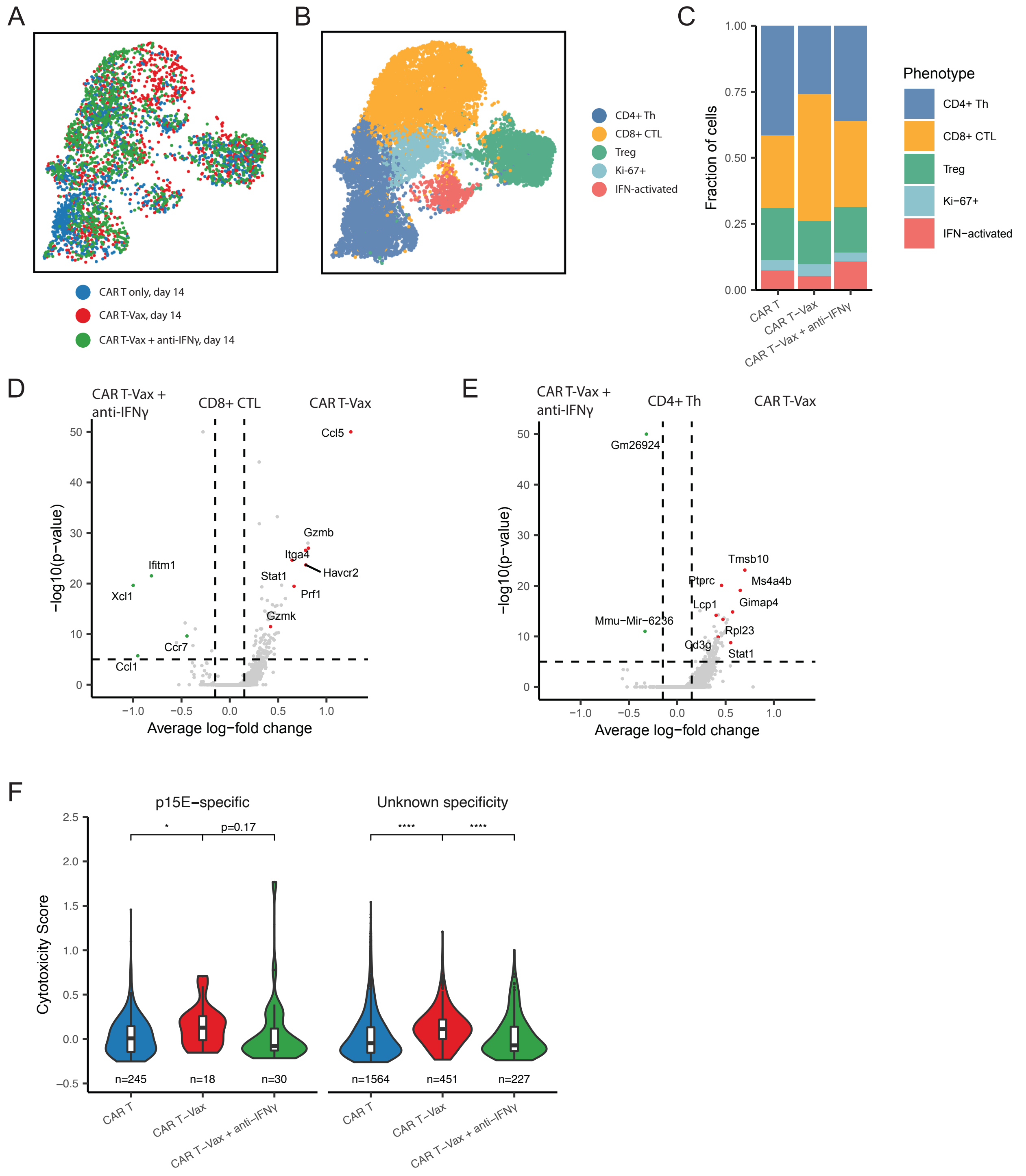


**Figure S5. Impact of IFN-γ blockade on the functionality and clonality of intratumoral endogenous T cells during CAR T-vax treatment at day 14.**

(A) UMAP of endogenous T cells obtained from the day 14 tumors in CAR-T- and CAR T-vax-treated mice in Fig. 2 as well as a group of mice receiving CAR T-vax + anti-IFNγ treatment. T cells were randomly down-sampled to show an even number of points from each treatment condition. T cells are colored by the treatment group.

(B) Curated clusters based on signature gene expression of day 14 T cells in Fig. S7A. Th, T helper cells. Treg, regulatory T cell. CTL, cytotoxic lymphocyte.
(C) Stacked charts showing proportions of different clusters within day 14 T cells under each treatment condition in Fig. S7A.
(D) Volcano plot showing differential gene expression in day 14 $CD8^+$ CTLs between CAR T-vax and CAR T-vax + anti IFNγ group in Fig. S7A.
(E) Volcano plot showing differential gene expression in day 14 $CD4^+$ Th cells between CAR T-vax and CAR T-vax + anti IFNγ group in Fig .S7A.
(F) Cytotoxicity score of endogenous retroviral antigen p15E-specific TILs and TILs of unknown specificity on day 14 from CAR-T, CAR T-vax and CAR T-vax + IFNγ groups. Data shown are mean ± 95% CI. ****, p<0.0001; *, p<0.05 by two-sided Wilcoxon rank-sum test. See methods for the definition and calculation of the cytotoxicity score.

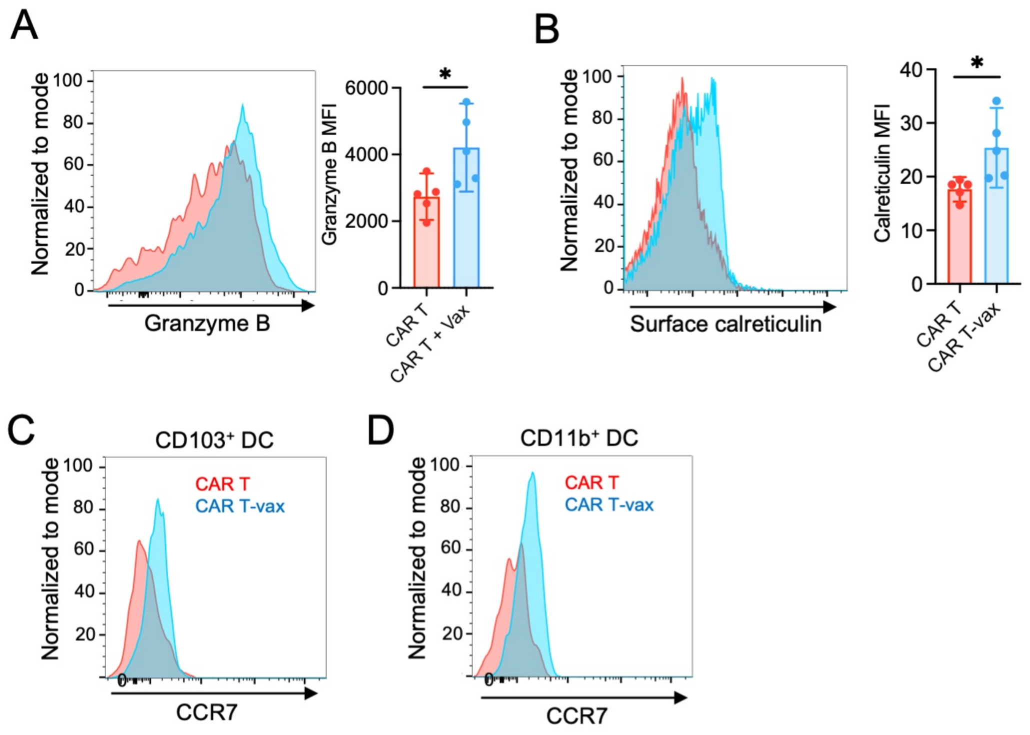


**Figure S6. Characterization of intratumoral CAR T, tumor cell, and DC responses to therapy.**
(A) Flow cytometry analysis showing intracellular Granzyme B expression in CAR T cells from mice in Fig. 6C.
(B)Tumors isolated from each group in Fig. 6C were dissociated into single cell suspensions for flow cytometry analysis of surface calreticulin expression on tumor cells in response to CAR-T vs. CAR T-vax therapy.
(C) Representative histogram showing surface CCR7 expression on intratumoral $CD103^+$ DCs from mice in Fig. 6E.
(D) Representative histogram showing surface CCR7 expression on intratumoral $CD11b^+$ DCs from mice in Fig. 6E.
Throughout, *n*=5 animals/group, Data shown are mean ± 95% CI. *, p<0.05 by Student's *t*-test.

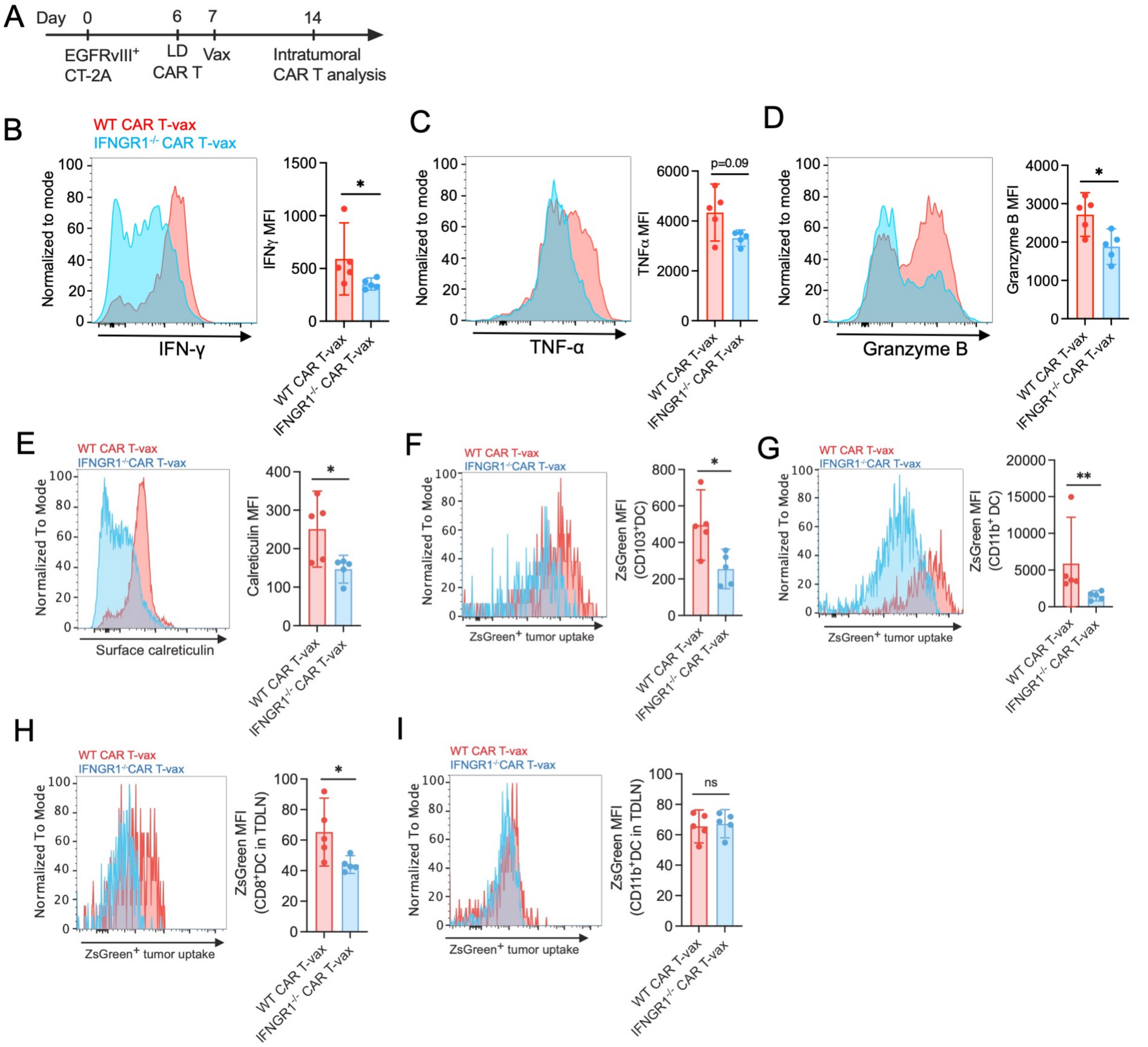


**Figure S7. Impact of CAR T-intrinsic IFN-γ signaling on CAR T polyfunctionality, subsequent tumor killing and stimulation of DC antigen uptake in response to vaccine boosting.**

(A-H) CD45.1$^{+}$ C57BL/6 mice bearing EGFRvIII$^{+}$CT-2A tumors received lymphodepletion (LD) and were treated with CD45.2$^{+}$ WT or IFNGR1$^{-/-}$CAR T-vax as shown in the timeline (A). Tumors isolated from each group at day 7 post CAR T-vax therapy were dissociated into single cell suspensions for flow cytometry analysis.

(B)IFN-γ expression in CD45.2$^{+}$ CAR T cells from WT or IFNGR1$^{-/-}$ CAR T-vax treated group.

(C)TNF-α expression in CD45.2$^{+}$ CAR T cells from WT or IFNGR1$^{-/-}$ CAR T-vax treated group.

(D) Granzyme B expression in CD45.2$^{+}$ CAR T cells from WT or IFNGR1$^{-/-}$ CAR T-vax treated group.

(E) Flow cytometry analysis showing surface expression of calreticulin on tumor cells.

(F) Flow cytometry analysis showing tumor antigen uptake by intratumoral CD45.2$^{+}$ CD103$^{+}$DCs. ZsGreen was used as a surrogate antigen in this experiment.

(G) Flow cytometry analysis showing tumor antigen uptake by intratumoral $CD45.2^+$ $CD11b^+$ DCs. ZsGreen was used as a surrogate antigen in this experiment.
(H) Flow cytometry analysis showing tumor antigen uptake by LN-resident $CD45.2^+$ $CD8^+$ DCs. ZsGreen was used as a surrogate antigen in this experiment.
(I) Flow cytometry analysis showing tumor antigen uptake by LN-resident $CD45.2^+$ $CD11b^+$ DCs. ZsGreen was used as a surrogate antigen in this experiment.
Throughout, a representative histogram from each treatment group and the summary data are shown. *n*=5 animals/group, data shown are mean ± 95% CI. **, $p<0.01$; *, $p<0.05$; ns, not significant by Student's *t*-test.

#### References

1. Moynihan, K. D. *et al.* Eradication of large established tumors in mice by combination immunotherapy that engages innate and adaptive immune responses. *Nature Medicine* 22, 1402–1410 (2016).

2. Ma, L. *et al.* Enhanced CAR–T cell activity against solid tumors by vaccine boosting through the chimeric receptor. *Science* 365, 162–168 (2019).

3. Clipstone, N. A. & Crabtree, G. R. Identification of calcineurin as a key signalling enzyme in T-lymphocyte activation. *Nature* 357, 695–697 (1992).

4. Liu, H. *et al.* Structure-based programming of lymph-node targeting in molecular vaccines. *Nature* 507, 519–522 (2014).

5. Kang, B. H. *et al.* Immunotherapy-induced antibodies to endogenous retroviral envelope glycoprotein confer tumor protection in mice. *Plos One* 16, e0248903 (2021).

6. Grace, B. E. *et al.* Identification of Highly Cross-Reactive Mimotopes for a Public T Cell Response in Murine Melanoma. *Front Immunol* 13, 886683 (2022).

7. Patro, R., Duggal, G., Love, M. I., Irizarry, R. A. & Kingsford, C. Salmon provides fast and bias-aware quantification of transcript expression. *Nat Methods* 14, 417–419 (2017).

8. Soneson, C., Love, M. I. & Robinson, M. D. Differential analyses for RNA-seq: transcript-level estimates improve gene-level inferences. *F1000research* 4, 1521 (2016).

9. Love, M. I., Huber, W. & Anders, S. Moderated estimation of fold change and dispersion for RNA-seq data with DESeq2. *Genome Biol* 15, 550 (2014).

10. Anders, S. & Huber, W. Differential expression analysis for sequence count data. *Genome Biol* 11, R106–R106 (2010).

11. Zhu, A., Ibrahim, J. G. & Love, M. I. Heavy-tailed prior distributions for sequence count data: removing the noise and preserving large differences. *Bioinformatics* 35, 2084–2092 (2019).

12. Mootha, V. K. *et al.* PGC-1α-responsive genes involved in oxidative phosphorylation are coordinately downregulated in human diabetes. *Nat Genet* 34, 267–273 (2003).

13. Subramanian, A. *et al.* Gene set enrichment analysis: A knowledge-based approach for interpreting genome-wide expression profiles. *Proc National Acad Sci* 102, 15545–15550 (2005).

14. Hughes, T. K. *et al.* Second-Strand Synthesis-Based Massively Parallel scRNA-Seq Reveals Cellular States and Molecular Features of Human Inflammatory Skin Pathologies. *Immunity* 53, 878-894.e7 (2020).

15. Stoeckius, M. *et al.* Cell Hashing with barcoded antibodies enables multiplexing and doublet detection for single cell genomics. *Genome Biol* 19, 224 (2018).

16. Macosko, E. Z. *et al.* Highly Parallel Genome-wide Expression Profiling of Individual Cells Using Nanoliter Droplets. *Cell* 161, 1202–1214 (2015).

17. Tu, A. A. *et al.* TCR sequencing paired with massively parallel 3′ RNA-seq reveals clonotypic T cell signatures. *Nat Immunol* 20, 1692–1699 (2019).

18. Gupta, N. T. *et al.* Change-O: a toolkit for analyzing large-scale B cell immunoglobulin repertoire sequencing data. *Bioinformatics* 31, 3356–3358 (2015).

19. Heiden, J. A. V. *et al.* pRESTO: a toolkit for processing high-throughput sequencing raw reads of lymphocyte receptor repertoires. *Bioinformatics* 30, 1930–1932 (2014).

20. Smith, T., Heger, A. & Sudbery, I. UMI-tools: modeling sequencing errors in Unique Molecular Identifiers to improve quantification accuracy. *Genome Res* 27, 491–499 (2017).

21. Charan, J. & Kantharia, N. D. How to calculate sample size in animal studies? *J Pharmacol Pharmacother* 4, 303–306 (2013).

22. Davila, M. L., Kloss, C. C., Gunset, G. & Sadelain, M. CD19 CAR-Targeted T Cells Induce Long-Term Remission and B Cell Aplasia in an Immunocompetent Mouse Model of B Cell Acute Lymphoblastic Leukemia. *PLOS ONE* 8, e61338-14 (2013).

23. Adachi, K. *et al.* IL-7 and CCL19 expression in CAR-T cells improves immune cell infiltration and CAR-T cell survival in the tumor. *Nature Biotechnology* 36, 346–351 (2018).
